# Interleukin-17 pathway activation in *Equus caballus* supporting limb laminitis

**DOI:** 10.1101/2020.04.27.063800

**Authors:** Lynne Cassimeris, Julie B. Engiles, Hannah Galantino-Homer

## Abstract

Supporting Limb Laminitis (SLL) is a painful and crippling secondary complication of orthopedic injuries and infections in horses, often resulting in euthanasia. Due to altered weight bearing, SLL causes structural alternations and inflammation of the interdigitating layers of specialized epidermal and dermal tissues, the lamellae, which suspend the equine distal phalanx from the hoof capsule. Activation of the interleukin-17 (IL-17)-dependent inflammatory pathway is an epidermal stress response that contributes to physiologic cutaneous wound healing as well as pathological skin conditions. To test the hypothesis that IL-17 pathway activation is involved in equine epidermal lamellae in SLL, we analyzed the expression of the IL-17 receptor subunit A and 11 genes upregulated by IL-17 in lamellar tissue isolated from Thoroughbreds euthanized due to naturally occurring SLL and in age and breed matched non-laminitic controls. The IL-17 Receptor A subunit was expressed in both non-laminitic and laminitic tissues. In severe acute SLL (n=7) compared to non-laminitic controls (n=8), quantitative PCR demonstrated ∼20-100 fold upregulation of ß defensin 4 (*E. caballus* gene *DEFB4B*) and *S100A9* genes. *DEFB4B* was also upregulated in developmental (n=8), moderate acute (n=7), and severe chronic (n=5) samples. By RT-PCR, *S100A8, MMP9*, and *PTSG2* (COX2) expression was upregulated in most or all severe acute SLL samples, whereas several other genes, *CCL2, CxCL8, TNFα, IL6* and *MMP1* were detected in some, but not all, severe acute samples. *PTGS2, CCL2, TNFα* and *IL6* were also expressed in some, but not all, developmental and moderate acute disease stages. Moreover, expression of *DEFB4* by in situ hybridization and calprotectin (S100A9/S100A8) protein by immunofluorescence was detected in keratinocytes, primarily in suprabasal cell layers, from SLL samples. These data support the hypothesis that the IL-17 inflammatory pathway is active in equine SLL, and that similarities exist between equine and human epidermal tissue responses to stresses and/or damage.

## Introduction

Equine laminitis is a common, progressive, crippling and currently incurable disease often necessitating euthanasia to end suffering from the painful loss of limb support. Healthy lamellar tissue connects the inner hoof wall to the distal phalanx (DP), forming a major component of the suspensory apparatus of the DP, a unique adaptation of equids that allows for suspension of almost the entire weight of the animal through the hoof capsule [1; 2]. The hoof is a modified epidermal appendage integrated into the musculoskeletal system by the epidermal and dermal lamellae, homologous to the nail bed, which are extensively folded to form primary and secondary lamellae to increase the surface area of attachment (Fig 1; [3,4]). Similar to skin, keratin 14 is expressed within lamellar epidermal basal cells [5-7]; however equine epidermal lamellae have several architectural and molecular modifications that differ from skin, including limited stratification into basal, suprabasal and cornified layers, and expression of unique keratins not found in equine or human skin [5; 6]. These modifications are thought to impart critical functional differences that allow equine lamellar tissue to bear significantly greater mechanical tensile and compressive forces compared to skin, consistent with the functional integration of the hoof capsule and lamellae into the musculoskeletal system [8].

**Fig 1.**
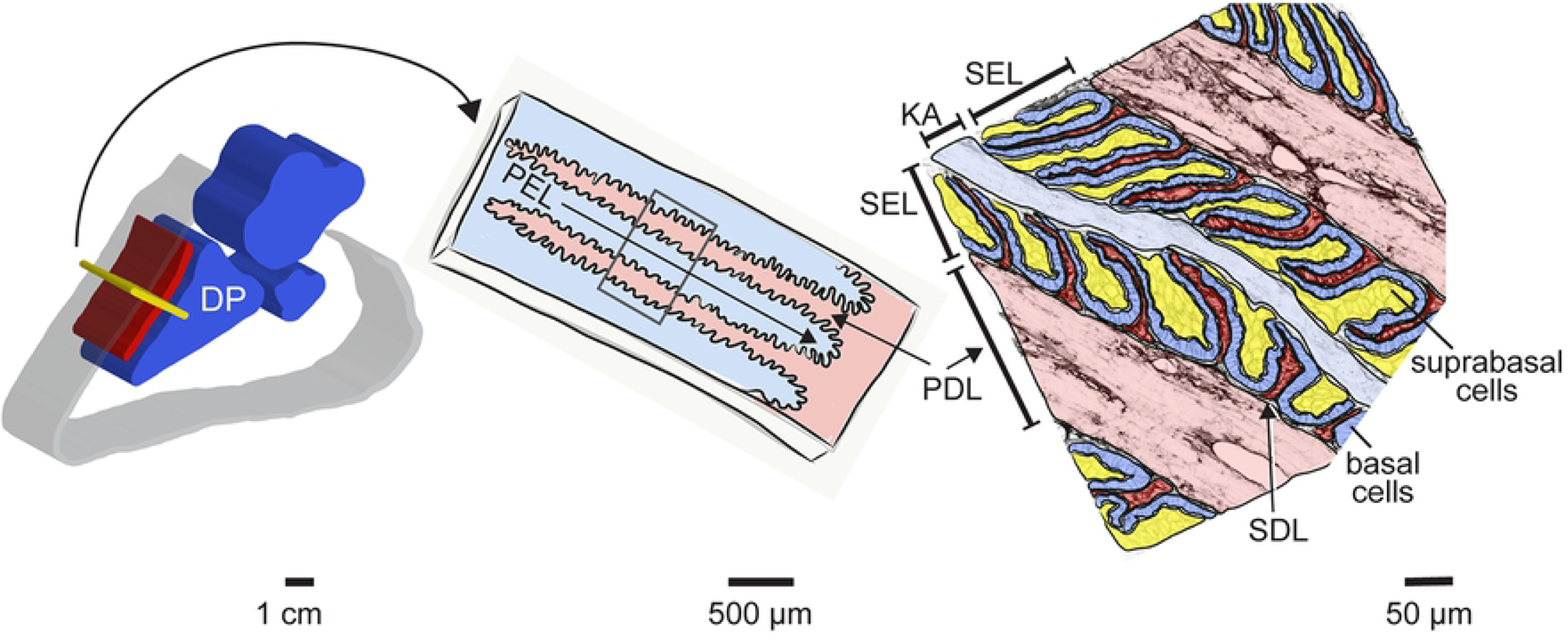
Schematic overview of equine hoof lamellar tissue anatomy. Left panel: cut away view of the midsagittal section of an equine foot outlining position of lamellar tissue (red) relative to hoof wall (grey) and distal phalanx (DP), middle phalanx, and distal sesamoid bones (blue). The transverse section plane used for tissue collection is shown in yellow. Center panel: a portion of the transverse section plane diagrammed schematically. The interdigitating primary epidermal lamellae (PEL) and primary dermal lamellae (PDL) are highlighted. Right panel: higher magnification illustration of lamellar tissue organization. Colors are overlaid on a tissue section stained to highlight cell membranes and extracellular matrix (rhodamine-tagged wheat germ agglutinin, see Methods). Darker blue overlays mark basal epidermal cells, yellow overlays mark suprabasal epidermal cells, light blue overlay highlights the keratinized axis (KA) of each PEL. The PEL consists of combined secondary epidermal lamellae (SEL) and KA. Dermal tissues are shown in red overlays, with light red highlighting PDL and darker red marking the secondary dermal lamellae (SDL).

In laminitis, equine epidermal lamellae become elongated, distorted and detached, leading to loss of mechanical support and failure to suspend the DP within the hoof capsule and, with progression to chronic laminitis, transform into an aberrantly thickened and dysplastic structure termed the “lamellar wedge” [2; 9-13]. There are three broad categories of laminitis associated with different etiologic risk factors: supporting limb laminitis (SLL), resulting from increased weight-bearing on the affected limb after pain from injury or bacterial infection reduces weight-bearing on a contralateral limb; endocrinopathic laminitis (EL) in which hyperinsulinemia is the key risk factor; and sepsis-related laminitis (SRL), triggered by systemic inflammatory conditions (reviewed by [13]). While differences in etiopathogenesis are reported for the different types of laminitis, several shared morphologic and molecular similarities exist, including keratinocyte apoptosis/necrosis; epidermal hyperplasia, acanthosis, dyskeratosis and hyperkeratosis; basement membrane separation; dermal inflammation; and progressive, deforming distal phalangeal osteolysis with medullary stromal activation and inflammation [9; 11; 14-19].

Interestingly, several histomorphologic features of equine laminitis are similar to those described for both physiologic cutaneous wound-healing and select chronic auto-inflammatory skin conditions, including human psoriasis. Human psoriasis is a chronic, heterogeneous immune-mediated inflammatory disease with genetic and epigenetic triggers that activate pro- inflammatory pathways, including IL-17A (referred to hereafter as IL-17) [18; 20; 21], resulting in cutaneous and nail manifestations including aberrant epidermal and subungual keratinocyte activation, differentiation and proliferation, vascular disturbances and inflammation, as well as extracutaneous manifestations [22]. A causal role for IL-17-dependent inflammation in driving psoriasis is highlighted by the success of monoclonal antibody-based therapies that bind IL-17 and block signaling through the IL-17 receptor [18; 21; 23]. Interestingly, a syndrome of psoriasis involving digits and nail beds, termed psoriatic onychodystrophy and psoriatic arthritis/dactylitis, includes lesions such as onycholysis, subungual acanthosis and hyperkeratosis, enthesitis and distal phalangeal osteolysis [24; 25]. Many of the changes listed above for human psoriasis have morphological similarities to equine laminitis and laminitis-associated distal phalangeal osteolysis [11]. IL-17 stimulated inflammation is also associated with a number of other human diseases, including rheumatoid arthritis, obesity-associated inflammation and cancer [19; 21; 26-29].

In human skin diseases and normal healing processes, both immune cells and keratinocytes respond to IL-17 directly via its receptor, a heterodimer of IL-17RA and IL-17RC (Fig 2; [18; 21; 30; 31]). Receptor activation results in increased expression of target genes either through enhanced translation or increased mRNA stability [21; 28; 33]. The cohort of target genes includes the ß defensins, with ß defensin 4 (human gene *DEF4B*) ranking as the most highly upregulated gene in human keratinocytes treated with IL-17 [34]. ß defensin 4 (also known as human ß defensin 2) is a biomarker for psoriasis and its decline provides a quantitative measure of immunotherapy efficacy [35; 36]. Additional effectors include cytokines, chemokines, antimicrobial proteins and matrix metalloproteinases [18; 21; 28; 33] as summarized in Fig 2.

**Fig 2.**
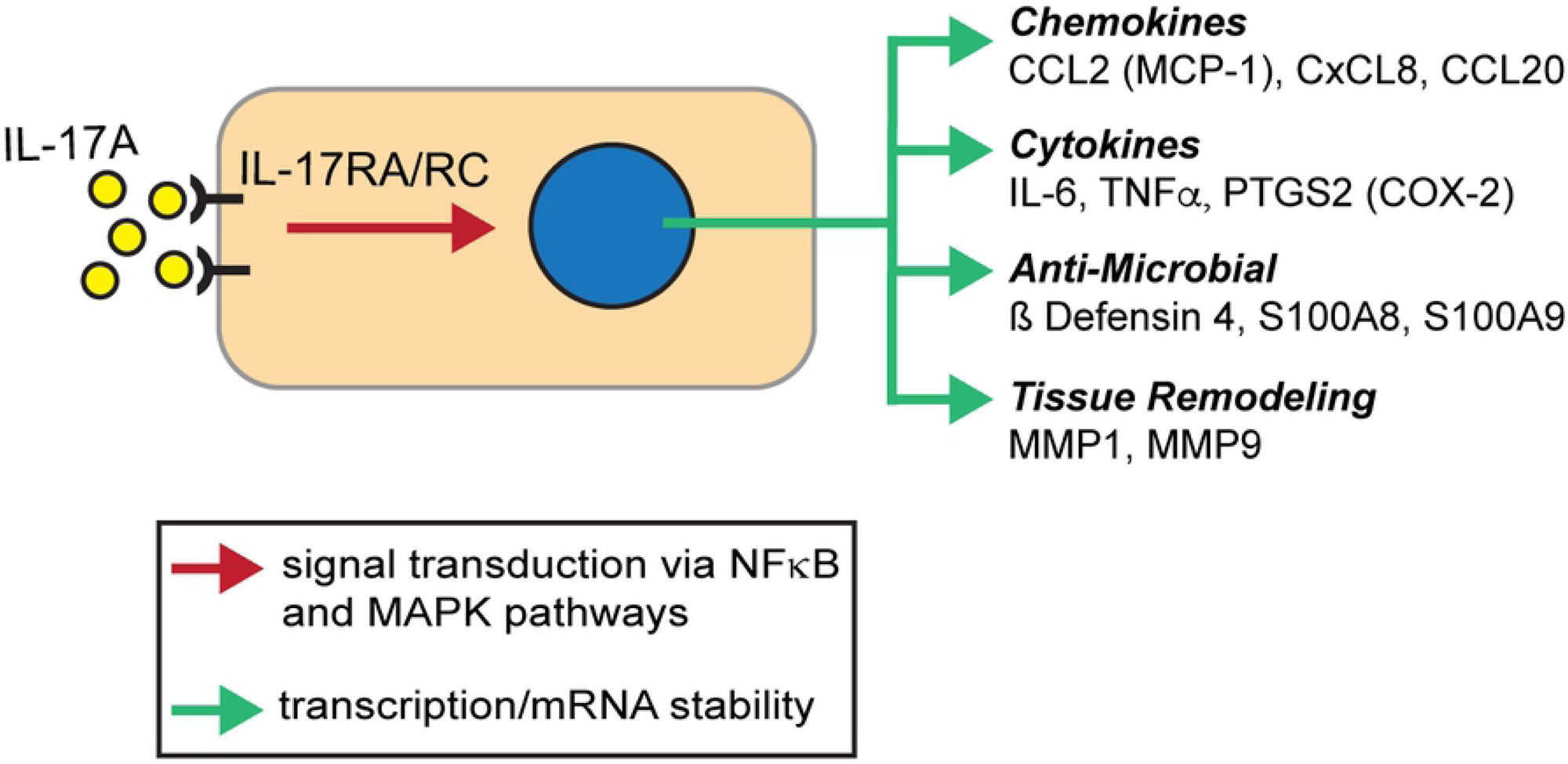
Overview of the IL-17 pathway and downstream target genes. IL-17 (IL-17A) binding to receptor heterodimer of IL-17RA and RC subunits activates downstream pathways to activate either new gene expression or increased mRNA stability for a cohort of target genes. The 11 genes assayed in this study are shown. See [21; 33; 37] for details on signal transduction.

Previous studies of inflammation in equine laminitis focused primarily on experimental models thought to mimic each of the three major laminitis categories (reviewed by [4; 13]). In general, most of these studies used treatments that mimic SRL, although several reports also examined models of EL or SLL [15; 38-49]. Models of SRL provide the most evidence for inflammation, either through upregulation of inflammatory mediators [15; 38; 39; 42-47] or leukocyte infiltration [50-52]. Recently, a model to mimic some aspects of SLL was reported [48]. Although this model did not result in clinical or histological laminitis and did not find upregulation of a similar set of inflammatory mediators identified in sepsis-related models, a relative increase in Hypoxia Inducible Factor (HIF)-1*α* protein levels in the supporting limb suggests that ischemia may contribute to pathogenesis of SLL [48]. Interestingly, a mouse model of experimental autoimmune encephalitis (EAE) demonstrated that hypoxia is a metabolic cue that can serve as a “triggering event” mediated through HIF-1*α* activation that initiates and perpetuates chronic inflammation and hypoxia through activation of IL-17 pathways that promote T_h_17 differentiation and inhibit T_reg_ differentiation [53]. Thus, variability or inconsistent upregulation of inflammatory mediators typically measured in different experimental models of equine laminitis described above raises the possibility that the degree or type of laminitis-associated inflammatory response may depend on the trigger used to induce the disease or that other inflammatory pathways not yet investigated may be involved [12].

Although experimental models of laminitis are important, examining specific inflammatory pathways in spontaneous cases of laminitis may provide additional information that helps elucidate critical components of the complex pathogenesis in this disease. Moreover, if mechanistic similarities exist between chronic inflammatory skin disease in humans and the development of chronic laminitis in horses, application of knowledge from human diseases could inform strategies for equine disease monitoring and/or treatment. To date, aside from the documentation of leukocyte infiltration, the presence of inflammatory mediators and pathway activation in natural (spontaneous) cases of laminitis has not been examined [4; 11; 50-52; 54; 55].

Here we test the hypothesis that the IL-17 pathway is active in natural cases of SLL using tissue from a cohort of young Thoroughbreds. Although natural cases provide a “temporal snap shot” of disease, typically at late or chronic stages where extensive damage has already occurred in the primarily affected foot, often horses euthanized for SLL demonstrate earlier, less severe stages of laminitis in other secondarily affected feet presumably due to related alterations in weight bearing [11]. From a cohort of archived lamellar tissues from horses euthanized for spontaneously occurring SLL that includes feet exhibiting varying stages and severity of disease, the expression of IL-17RA and eleven IL-17 target genes were examined by either RT-PCR or qPCR. Cellular localization was examined for two effectors by in situ hybridization (ß defensin 4; *E. caballus* gene name *DEFB4B*) or immunofluorescence (calprotectin, a dimer of S100A8/S100A9 proteins) to ask whether the keratinocytes themselves respond by upregulating expression of IL-17 target genes.

## Methods

### Ethics statement

The archived equine tissue samples used in this study were collected under protocols titled ‘Pathophysiology of Equine Laminitis (By-products only)’ and ‘Equine Laminitis Tissue Bank’. These protocols were approved by the University of Pennsylvania Institutional Animal Care and Use Committee (protocol #801950 and #804262, respectively). Euthanasia of the horses was carried out in accordance with the recommendations in the Guide for the Care and Use of Agricultural Animals in Research in Teaching Federation for Animal Science Societies) and the AVMA Guidelines for the Euthanasia of Animals (American Veterinary Medical Association) by overdose with pentobarbital sodium and phenytoin sodium.

### Subjects and tissue retrieval

Samples were obtained from the Laminitis Discovery Database, a tissue repository housed in the Galantino-Homer laboratory at the University of Pennsylvania School of Veterinary Medicine, New Bolton Center [56]. The cohort included 14 Thoroughbred SLL cases of reduced-weight bearing lameness with clinical signs and gross and/or histological lesions consistent with laminitis in at least one limb and 7 age-matched Thoroughbred controls. At the time of euthanasia, all control and SLL case Thoroughbreds were currently in, or recently retired due to injury, from racing or training to race, aged 2-7 years, and included females, males and castrated males, as listed in Table 1. All control horses and one SLL case were donated for research and teaching; the remaining SLL cases were client-owned animals submitted for necropsy following euthanasia due to SLL. Lamellar tissue, including epidermal and dermal lamellae, was collected immediately after euthanasia as previously described [7; 11; 56; 57]. Tissue samples were immediately either snap frozen in liquid nitrogen and stored in liquid nitrogen until processed for RNA extraction, or formalin-fixed/paraffin-embedded (FFPE) and sectioned for microscopy experiments [7; 57; 58]. Tissue sections were oriented transverse to the axis of the limb and include the entire abaxial, middle, and axial regions (relative to the DP) of the primary epidermal lamellae as well as the sub-lamellar dermal tissue located between the DP and epidermal lamellae (see Fig 1).

**Table 1.**
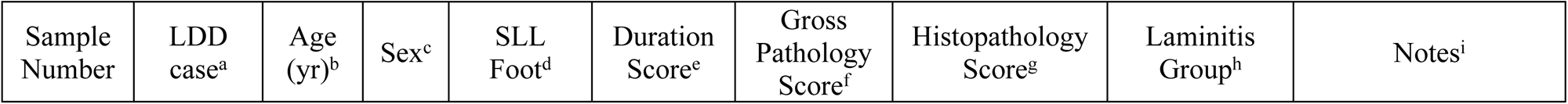
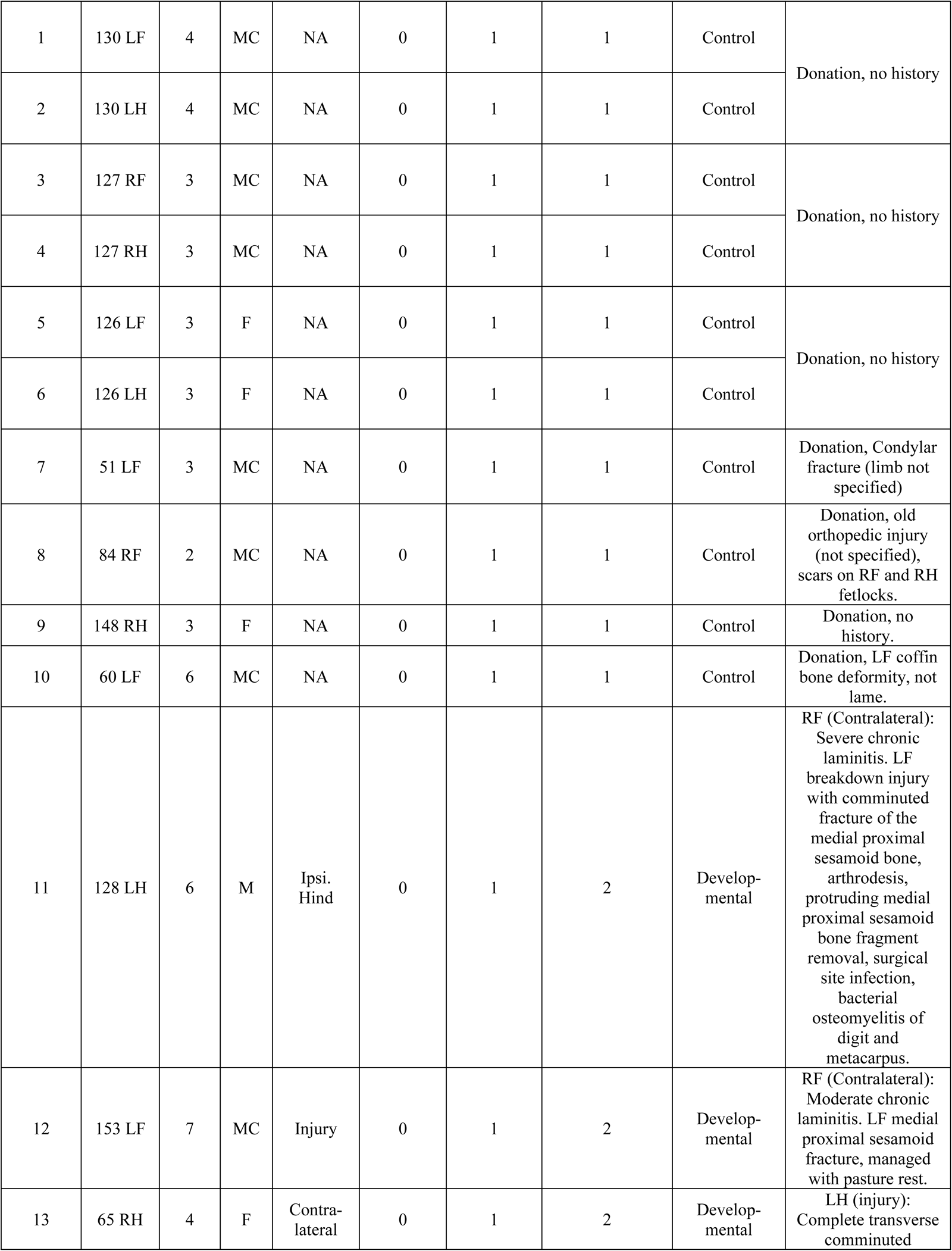
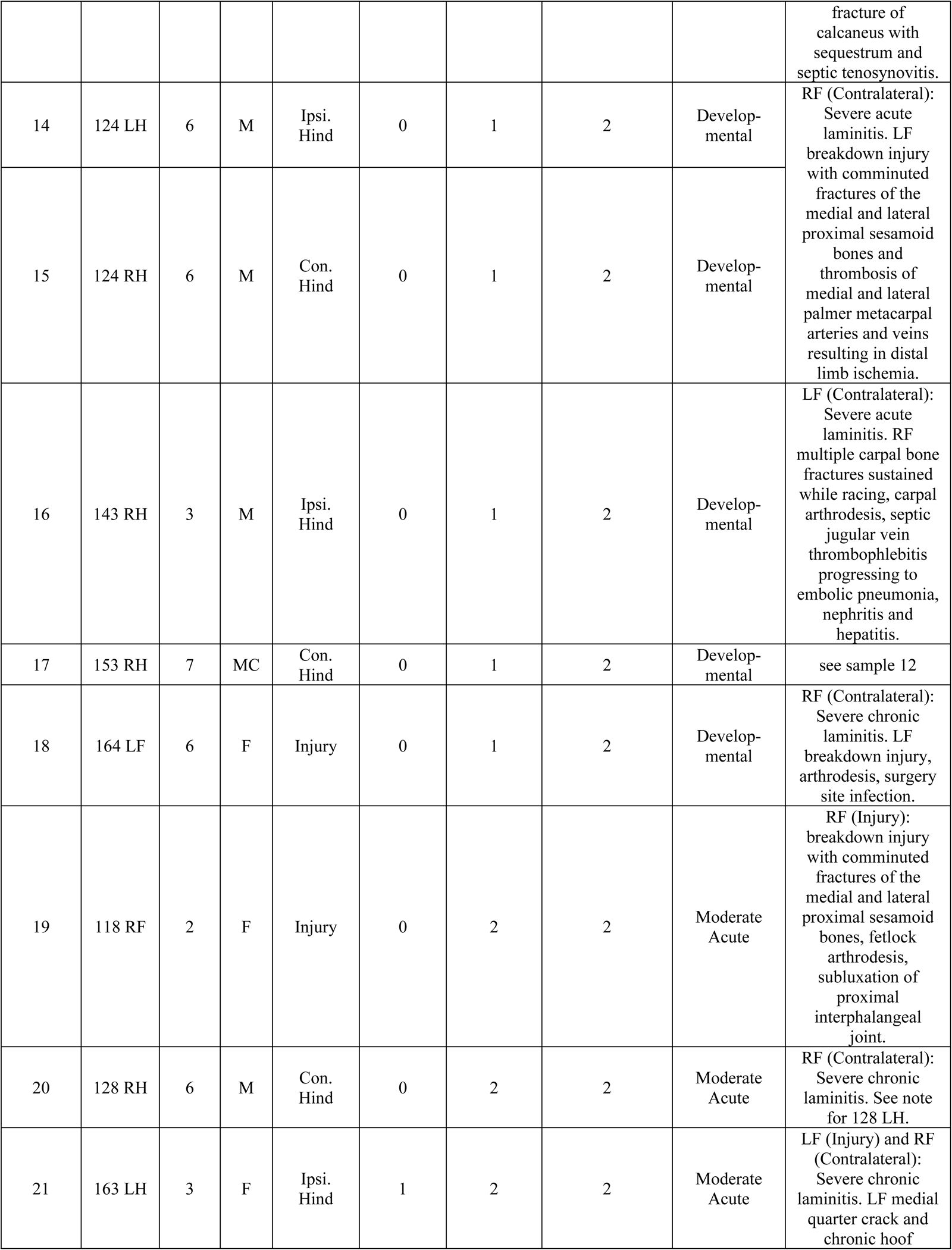
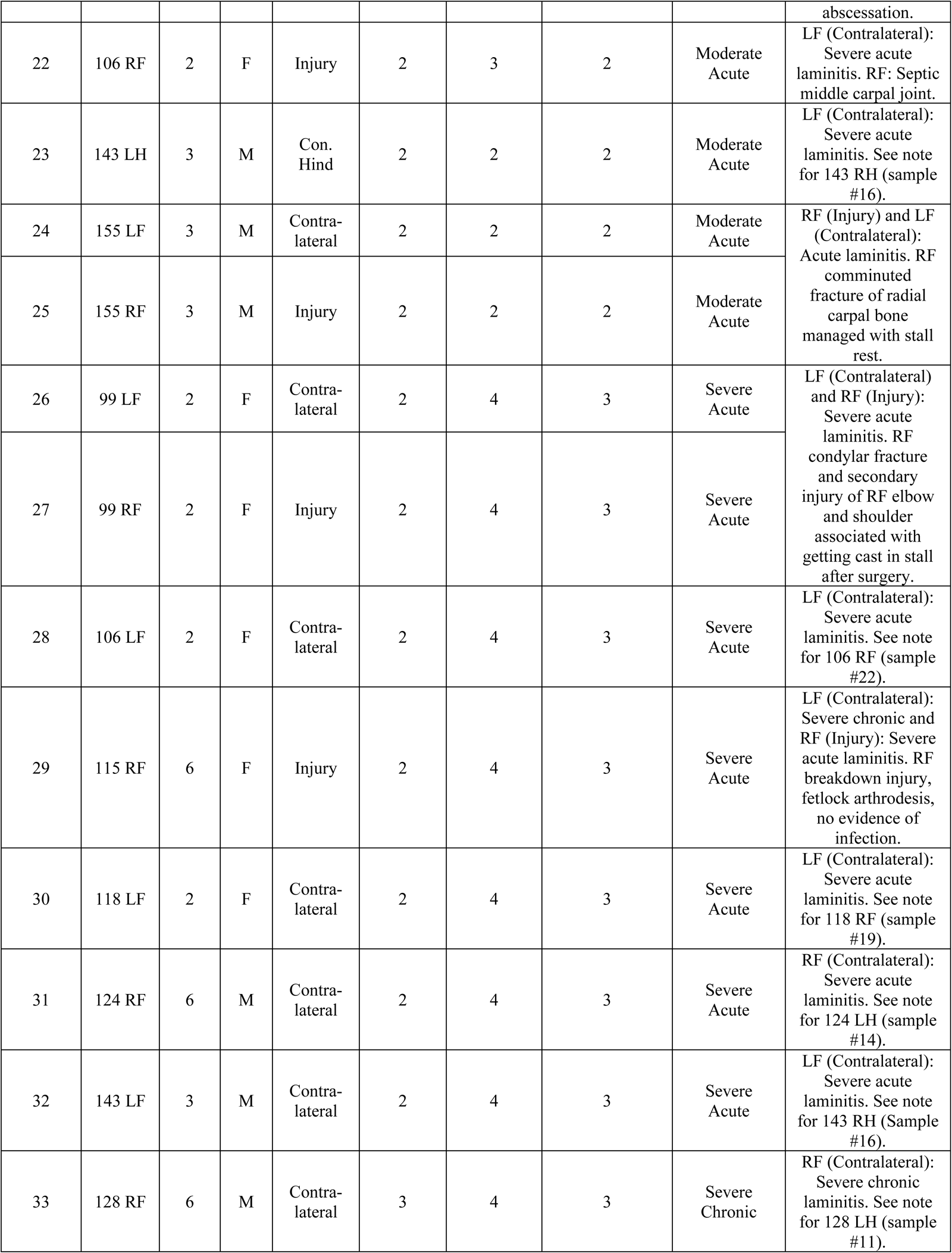
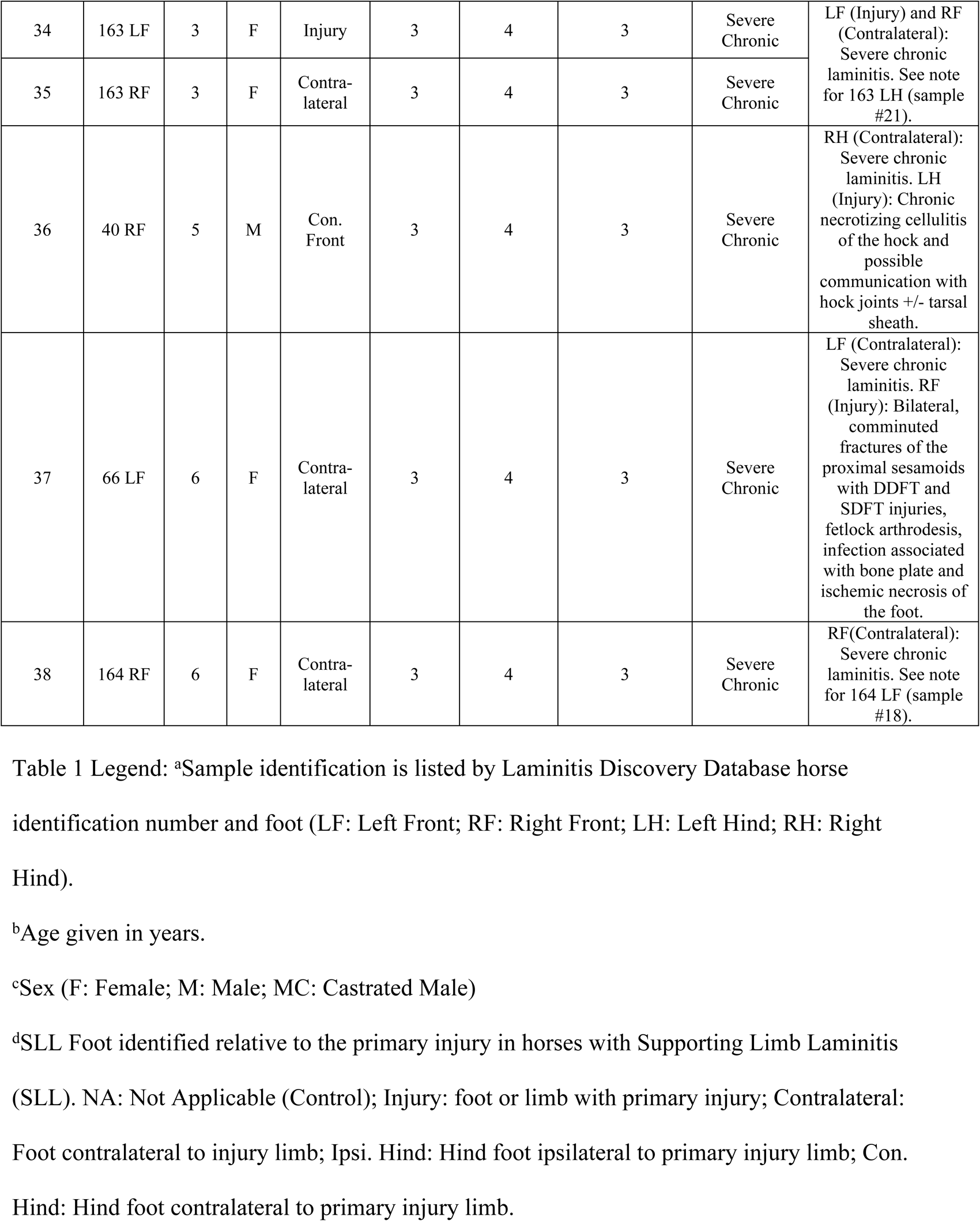
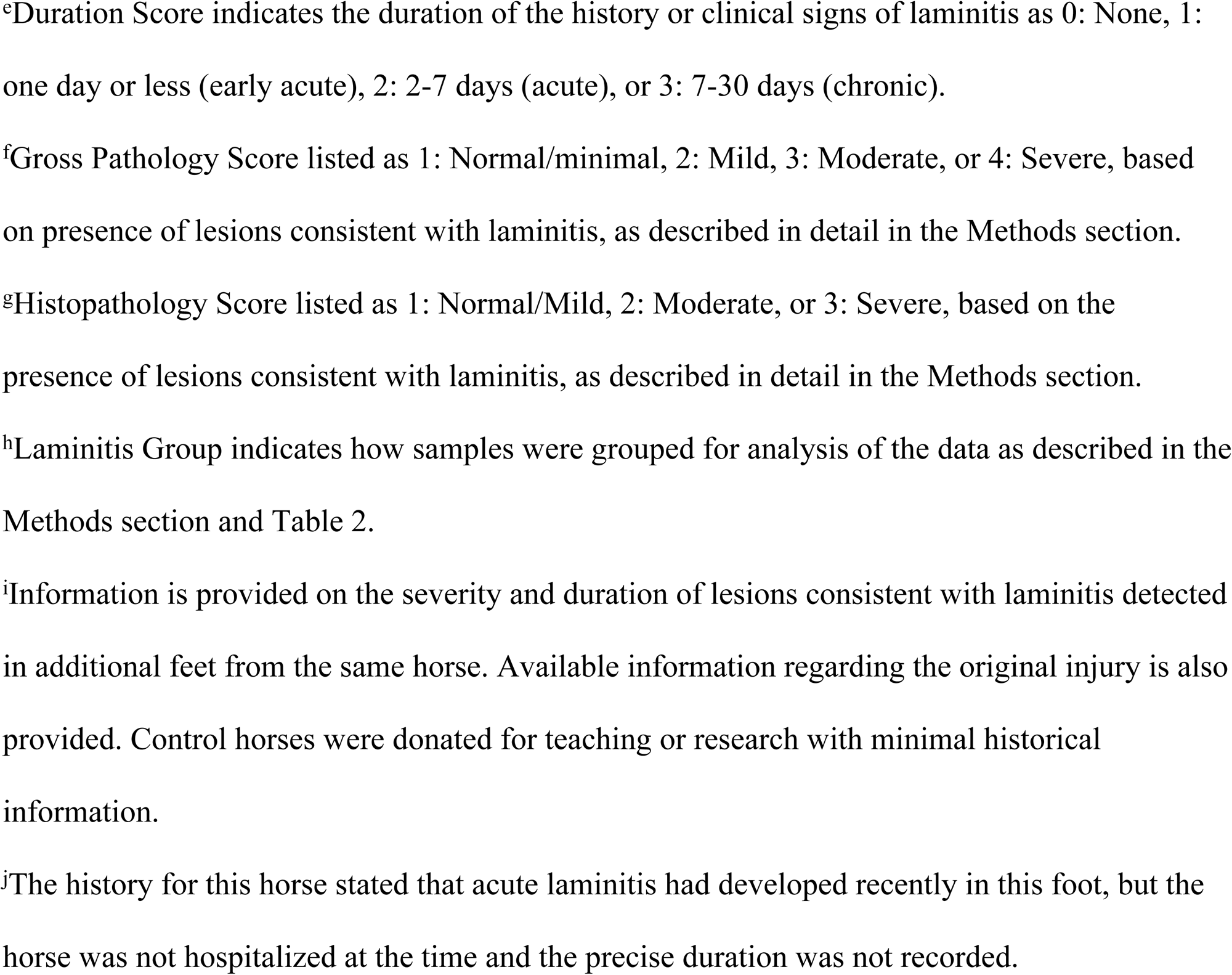
Sample summary of signalment, duration of clinical signs of laminitis, gross pathology and histopathology for lamellar tissue from Thoroughbred horses used for RNA extraction.

Legend: <sup>a</sup>Sample identification is listed by Laminitis Discovery Database horse identification number and foot (LF: Left Front; RF: Right Front; LH: Left Hind; RH: Right Hind).
| Sample Number | LDD case<sup>a</sup> | Age (yr)<sup>b</sup> | Sex<sup>c</sup> | SLL Foot<sup>d</sup> | Duration Score<sup>e</sup> | Gross Pathology Score<sup>f</sup> | Histopathology Score<sup>g</sup> | Laminitis Group<sup>h</sup> | Notes<sup>i</sup> |
| --- | --- | --- | --- | --- | --- | --- | --- | --- | --- |
| 1 | 130 LF | 4 | MC | NA | 0 | 1 | 1 | Control | Donation, no history |
| 2 | 130 LH | 4 | MC | NA | 0 | 1 | 1 | Control |  |
| 3 | 127 RF | 3 | MC | NA | 0 | 1 | 1 | Control | Donation, no history |
| 4 | 127 RH | 3 | MC | NA | 0 | 1 | 1 | Control |  |
| 5 | 126 LF | 3 | F | NA | 0 | 1 | 1 | Control | Donation, no history |
| 6 | 126 LH | 3 | F | NA | 0 | 1 | 1 | Control |  |
| 7 | 51 LF | 3 | MC | NA | 0 | 1 | 1 | Control | Donation, Condylar fracture (limb not specified) |
| 8 | 84 RF | 2 | MC | NA | 0 | 1 | 1 | Control | Donation, old orthopedic injury (not specified), scars on RF and RH fetlocks. |
| 9 | 148 RH | 3 | F | NA | 0 | 1 | 1 | Control | Donation, no history. |
| 10 | 60 LF | 6 | MC | NA | 0 | 1 | 1 | Control | Donation, LF coffin bone deformity, not lame. |
| 11 | 128 LH | 6 | M | Ipsi. Hind | 0 | 1 | 2 | Developmental | RF (Contralateral): Severe chronic laminitis. LF breakdown injury with comminuted fracture of the medial proximal sesamoid bone, arthrodesis, protruding medial proximal sesamoid bone fragment removal, surgical site infection, bacterial osteomyelitis of digit and metacarpus. |
| 12 | 153 LF | 7 | MC | Injury | 0 | 1 | 2 | Developmental | RF (Contralateral): Moderate chronic laminitis. LF medial proximal sesamoid fracture, managed with pasture rest. |
| 13 | 65 RH | 4 | F | Contralateral | 0 | 1 | 2 | Developmental | LH (injury): Complete transverse comminuted fracture of calcaneus with sequestrum and septic tenosynovitis. |
| 14 | 124 LH | 6 | M | Ipsi. Hind | 0 | 1 | 2 | Developmental | RF (Contralateral): Severe acute laminitis. LF breakdown injury with comminuted fractures of the medial and lateral proximal sesamoid bones and thrombosis of medial and lateral palmar metacarpal arteries and veins resulting in distal limb ischemia. |
| 15 | 124 RH | 6 | M | Con. Hind | 0 | 1 | 2 | Developmental |  |
| 16 | 143 RH | 3 | M | Ipsi. Hind | 0 | 1 | 2 | Developmental | LF (Contralateral): Severe acute laminitis. RF multiple carpal bone fractures sustained while racing, carpal arthrodesis, septic jugular vein thrombophlebitis progressing to embolic pneumonia, nephritis and hepatitis. |
| 17 | 153 RH | 7 | MC | Con. Hind | 0 | 1 | 2 | Developmental | see sample 12 |
| 18 | 164 LF | 6 | F | Injury | 0 | 1 | 2 | Developmental | RF (Contralateral): Severe chronic laminitis. LF breakdown injury, arthrodesis, surgery site infection. |
| 19 | 118 RF | 2 | F | Injury | 0 | 2 | 2 | Moderate Acute | RF (Injury): breakdown injury with comminuted fractures of the medial and lateral proximal sesamoid bones, fetlock arthrodesis, subluxation of proximal interphalangeal joint. |
| 20 | 128 RH | 6 | M | Con. Hind | 0 | 2 | 2 | Moderate Acute | RF (Contralateral): Severe chronic laminitis. See note for 128 LH. |
| 21 | 163 LH | 3 | F | Ipsi. Hind | 1 | 2 | 2 | Moderate Acute | LF (Injury) and RF (Contralateral): Severe chronic laminitis. LF medial quarter crack and chronic hoof abscessation. |
| 22 | 106 RF | 2 | F | Injury | 2 | 3 | 2 | Moderate Acute | LF (Contralateral): Severe acute laminitis. RF: Septic middle carpal joint. |
| 23 | 143 LH | 3 | M | Con. Hind | 2 | 2 | 2 | Moderate Acute | LF (Contralateral): Severe acute laminitis. See note for 143 RH (sample #16). |
| 24 | 155 LF | 3 | M | Contra-lateral | 2 | 2 | 2 | Moderate Acute | RF (Injury) and LF (Contralateral): Acute laminitis. RF comminuted fracture of radial carpal bone managed with stall rest. |
| 25 | 155 RF | 3 | M | Injury | 2 | 2 | 2 | Moderate Acute |  |
| 26 | 99 LF | 2 | F | Contra-lateral | 2 | 4 | 3 | Severe Acute | LF (Contralateral) and RF (Injury): Severe acute laminitis. RF condylar fracture and secondary injury of RF elbow and shoulder associated with getting cast in stall after surgery. |
| 27 | 99 RF | 2 | F | Injury | 2 | 4 | 3 | Severe Acute |  |
| 28 | 106 LF | 2 | F | Contra-lateral | 2 | 4 | 3 | Severe Acute | LF (Contralateral): Severe acute laminitis. See note for 106 RF (sample #22). |
| 29 | 115 RF | 6 | F | Injury | 2 | 4 | 3 | Severe Acute | LF (Contralateral): Severe chronic and RF (Injury): Severe acute laminitis. RF breakdown injury, fetlock arthrodesis, no evidence of infection. |
| 30 | 118 LF | 2 | F | Contra-lateral | 2 | 4 | 3 | Severe Acute | LF (Contralateral): Severe acute laminitis. See note for 118 RF (sample #19). |
| 31 | 124 RF | 6 | M | Contra-lateral | 2 | 4 | 3 | Severe Acute | RF (Contralateral): Severe acute laminitis. See note for 124 LH (sample #14). |
| 32 | 143 LF | 3 | M | Contra-lateral | 2 | 4 | 3 | Severe Acute | LF (Contralateral): Severe acute laminitis. See note for 143 RH (Sample #16). |
| 33 | 128 RF | 6 | M | Contra-lateral | 3 | 4 | 3 | Severe Chronic | RF (Contralateral): Severe chronic laminitis. See note for 128 LH (sample #11). |
| 34 | 163 LF | 3 | F | Injury | 3 | 4 | 3 | Severe Chronic | LF (Injury) and RF (Contralateral): Severe chronic laminitis. See note for 163 LH (sample #21). |
| 35 | 163 RF | 3 | F | Contra-lateral | 3 | 4 | 3 | Severe Chronic |  |
| 36 | 40 RF | 5 | M | Con. Front | 3 | 4 | 3 | Severe Chronic | RH (Contralateral): Severe chronic laminitis. LH (Injury): Chronic necrotizing cellulitis of the hock and possible communication with hock joints +/- tarsal sheath. |
| 37 | 66 LF | 6 | F | Contra-lateral | 3 | 4 | 3 | Severe Chronic | LF (Contralateral): Severe chronic laminitis. RF (Injury): Bilateral, comminuted fractures of the proximal sesamoids with DDFT and SDFT injuries, fetlock arthrodesis, infection associated with bone plate and ischemic necrosis of the foot. |
| 38 | 164 RF | 6 | F | Contra-lateral | 3 | 4 | 3 | Severe Chronic | RF(Contralateral): Severe chronic laminitis. See note for 164 LF (sample #18). |

<sup>e</sup>Duration Score indicates the duration of the history or clinical signs of laminitis as 0: None, 1: one day or less (early acute), 2: 2-7 days (acute), or 3: 7-30 days (chronic).
<sup>f</sup>Gross Pathology Score listed as 1: Normal/minimal, 2: Mild, 3: Moderate, or 4: Severe, based on presence of lesions consistent with laminitis, as described in detail in the Methods section.
<sup>g</sup>Histopathology Score listed as 1: Normal/Mild, 2: Moderate, or 3: Severe, based on the presence of lesions consistent with laminitis, as described in detail in the Methods section.
<sup>h</sup>Laminitis Group indicates how samples were grouped for analysis of the data as described in the Methods section and Table 2.
<sup>i</sup>Information is provided on the severity and duration of lesions consistent with laminitis detected in additional feet from the same horse. Available information regarding the original injury is also provided. Control horses were donated for teaching or research with minimal historical information.
<sup>j</sup>The history for this horse stated that acute laminitis had developed recently in this foot, but the horse was not hospitalized at the time and the precise duration was not recorded.

SLL cases were defined by the presence of reduced weight-bearing lameness due to injury or bacterial infection affecting one limb and the clinical expression of signs compatible with SLL in at least one other limb at the time of euthanasia, as reported by the attending veterinarian, and confirmed by gross pathology and histopathology evaluations as described below. The 7 age-matched controls were euthanized primarily for lameness due to non-laminitic orthopedic disease and showed few gross or histological lesions compatible with laminitis (Table 1).

### Antemortem Clinical Characterization and Postmortem Gross/Histological Evaluation

Each foot was scored for antemortem presence and duration (in days) of clinical signs compatible with laminitis, as well as gross and histologic postmortem lesions compatible with non-laminitic control and 4 SLL categories (developmental, moderate acute, severe acute, or severe chronic), as follows and as summarized in Table 2: **Non-laminitic**: No clinical signs of laminitis, a normal or minimal gross pathology score (gross pathology score 1) and no or minimal histological lesions compatible with laminitis (histopathology score 1); **Developmental laminitis**: No clinical signs of laminitis, a normal or minimal gross pathology score (gross pathology score 1), but histological lesions compatible with moderate lamellar damage are present (histopathology score 2); **Moderate acute laminitis**: 1-7 days duration of active clinical signs of laminitis, moderate gross pathology lesions (gross pathology score of 2-3), and histological lesions compatible with moderate laminitis (histopathology score 2); **Severe acute laminitis**: 1-7 days duration of active clinical signs of laminitis, severe gross pathology lesions (gross pathology score of 4), and histological lesions compatible with severe laminitis (histopathology score 3); **Severe chronic laminitis**: > 7 days duration of active clinical signs of laminitis, severe gross pathology lesions (gross pathology score of 4), and histological lesions compatible with severe laminitis (histopathology score 3).

**Table 2.** Laminitis group parameters.

| Laminitis Group | Clinical Duration Score<sup>a</sup> | Gross Pathology Score<sup>b</sup> | Histopathology Score<sup>c</sup> |
| --- | --- | --- | --- |
| Control | 0 | 1 | 1 |
| Developmental | 0 | 1 | 1 |
| Moderate Acute | 0-2 | 2-3 | 2 |
| Severe Acute | 2 | 4 | 3 |
| Severe Chronic | 3 | 4 | 3 |

<sup>a</sup>Duration Score indicates the duration of the history or clinical signs of laminitis as 0: None, 1: one day or less (early acute), 2: 2-7 days (acute), or 3: 7-30 days (chronic).
<sup>b</sup>Gross Pathology Score listed as 1: Normal/minimal, 2: Mild, 3: Moderate, or 4: Severe, based on presence of lesions consistent with laminitis, as previously described [Cassimeris, '19].
<sup>c</sup>Histopathology Score listed as 1: Normal/Mild, 2: Moderate, or 3: Severe, based on the presence of lesions consistent with laminitis, as previously described [59].

Gross pathology was assessed from direct observations and photographic images captured immediately after mid-sagittal sectioning of the foot and was scored to group SLL limbs into 4 severity categories (1: minimal or no lesions; 2: mild lesions; 3: moderate lesions; 4 severe lesions) based on the presence of lesions as previously described [59]). Quantitative and qualitative histopathology was assessed on FFPE mid-dorsal lamellar tissue samples that were sectioned and stained with hematoxylin and eosin stain and with Periodic acid-Schiff (PAS)- hematoxylin (PASH) stain, as described previously [59]. The detailed histopathology scoring and analysis for the sample set used for this study will be published separately. The summary histopathology scores presented in Table 1 and Table 2 represent: 1) Normal/Mild, none or focal distribution of lesions compatible with laminitis, 2) Moderate, multifocal to regional distribution of lesions compatible with laminitis, or 3) Severe, global distribution of lesions compatible with laminitis. The evaluation of histopathology lesions was performed as previously described [59].

### Oligonucleotide primers

Primers were designed to amplify equine sequences encoding regions of IL-17RA (receptor subunit A), as well as 11 products of the IL-17 pathway (Table 3). Primer sequences for several of the target genes were based on previous publications (Table 3). Additional primers were designed using tools available through either NCBI (https://www.ncbi.nlm.nih.gov/tools/primer-blast/) or OligoPerfect Primer Designer tool (ThermoFisher Scientific; https://www.thermofisher.com/us/en/home/life-science/oligonucleotides-primers-probes-genes/custom-dna-oligos/oligo-design-tools/oligoperfect.html). *RACK1* (formerly named *GNB2L1*) was used as the reference gene for PCR-based experiments based on previous work demonstrating stable expression of this gene in lamellar tissue from horses with experimental laminitis and non-laminitic controls [60]. The *RACK1* primers amplify a region spanning an intron/exon border and amplify a 200 bp sequence from cDNA and a 741 bp sequence from genomic DNA, providing confirmation that all cDNA samples were free of genomic DNA contamination. With the exception of *IL6* and *PTGS2*, primers for target genes also spanned more than one exon, also confirming that experiments identified fully processed mRNAs. Oligonucleotide primers were synthesized by Integrated DNA Technologies, Inc. (Coralville, IA, USA).

**Table 3.**
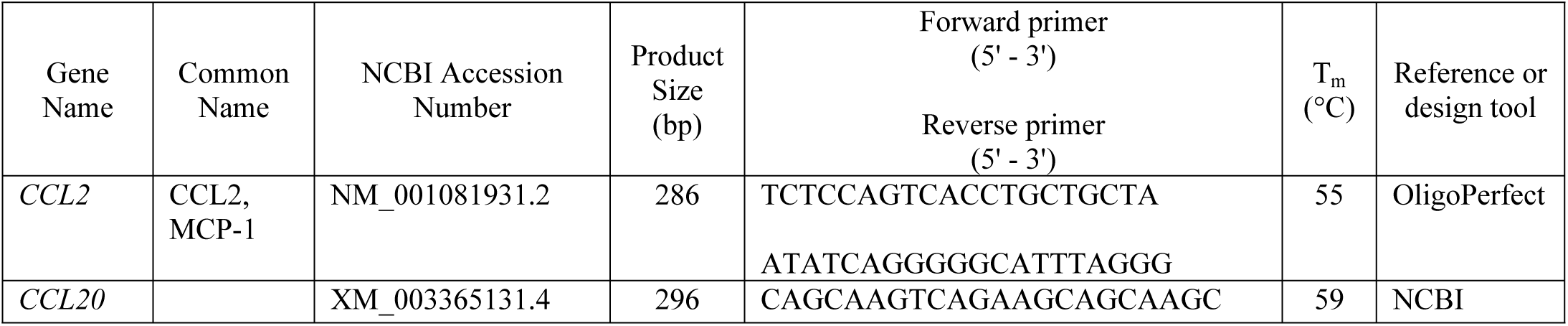
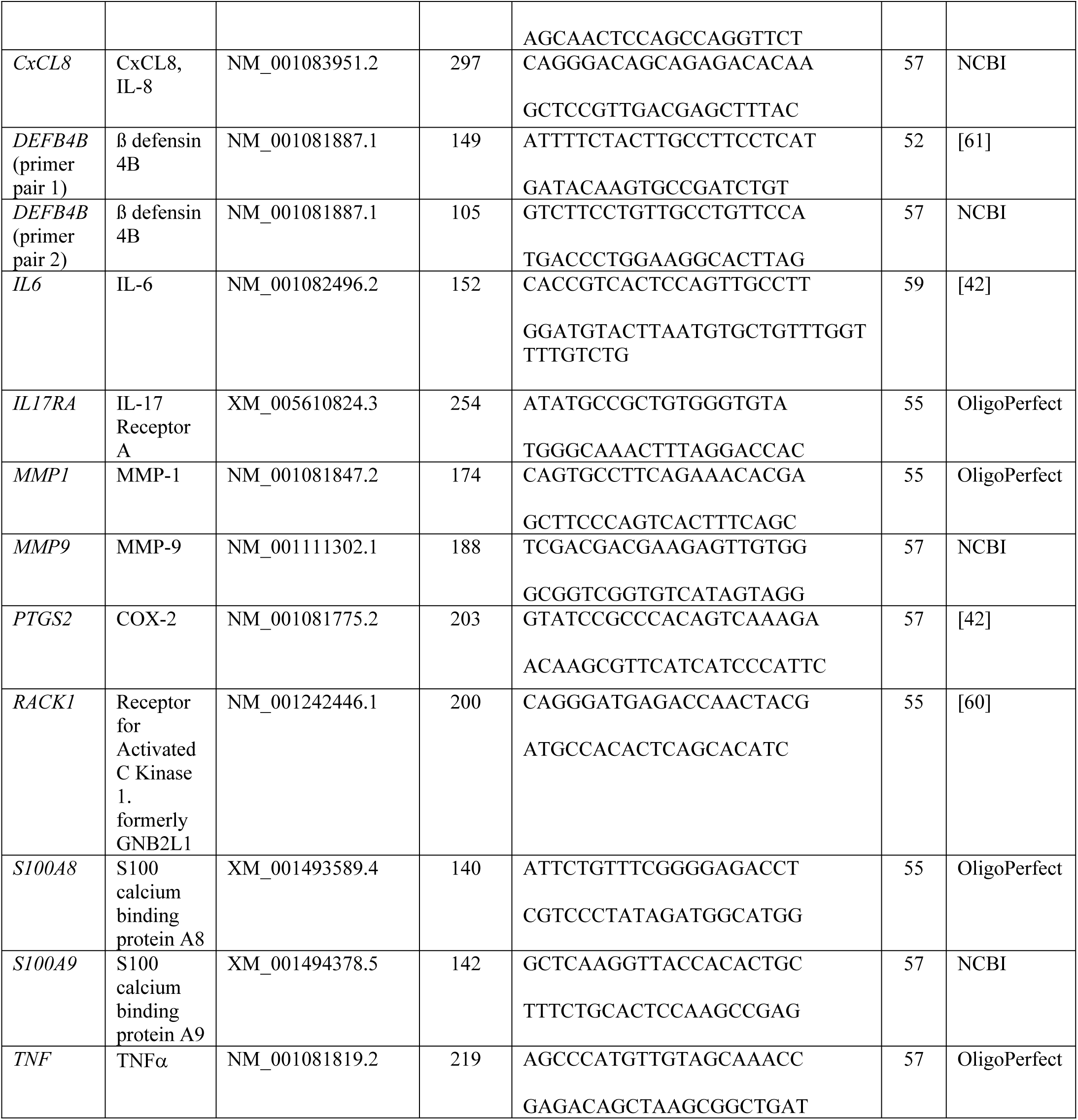
Equine PCR Primer Sequences. Gene names are as given for the NCBI accession numbers. Melting temperatures (T_m_) were provided by Integrated DNA Technologies.

### RNA extraction and cDNA synthesis

Archived snap-frozen tissue was pulverized and RNA extracted as described in detail previously [7]. Briefly, total RNA was extracted from pulverized tissue using the RNeasy Fibrous Tissue Mini Kit (Qiagen, Valencia, CA, USA) with modifications from the manufacturer’s instructions necessary for the highly fibrous lamellar tissue. Pulverized tissue samples were vortexed gently in 900 µl of Buffer RLT and proteinase K, incubated at 55°C for 10 min and homogenized by passing 5-10x through an 18 g hypodermic needle and syringe. The resulting material was clarified at 10,000 x g for 10 min. The supernatant was combined with 0.5 volume of 100% ethanol and the solution applied to the RNeasy Mini column in 700 µl increments, and then spun at 10,000 x g. RNA was eluted from the RNeasy column by adding 350 µl Buffer RW1 and centrifugation for 15 s at 10,000 x g. DNase treatment and subsequent buffer RW1 and RPE steps were then performed according to the manufacturer’s instructions. RNA was eluted in a total volume of 50 µl of RNase-free water. Total RNA was quantified using a Nanodrop 2000 spectrophotometer (Thermo Fisher Scientific, Waltham, MA, USA). The quality of the RNA preparations was confirmed by A260/A280 ratios of 2.0 ± 0.1 for all samples used here.

cDNAs were prepared from each RNA sample using a TaqManTM reverse transcription kit using random hexamer primers (ThermoFisher Scientific, catalog number N8080234) according the the manufacturer’s instructions. Total reaction volumes were 20 - 40 µl and included 100 ng of RNA per each 20 µL volume. Reverse transcription was carried out for 30 min at 37°C.

### RT-PCR

PCR amplification reactions of 25 µL total volume included ThermoPol reaction buffer, 10 ng cDNA template, 0.625 units Taq DNA polymerase (New England Biolabs), 0.2 µM forward and reverse primers (Table 3), and 200 µM dNTPs. Amplification reactions were run using an Eppendorf Masterflex thermocycler. Amplimers were detected by electrophoresis on 2% agarose (Sigma-Aldrich) gels and visualized with ethidium bromide (MP Biomedicals LLC., Irvine, CA). Gels were imaged using a BioRad ChemiDoc MP system running Image Lab 5.1 software and exported as TIFF files. The black/white scale was inverted in Photoshop (version CC 2018; Adobe) for clarity.

### qPCR

The qPCR amplification reactions totaled 20 µL that included Power SYBR Green PCR Master Mix (Applied Biosystems), 0.5 µM forward and reverse primers, and 5 ng template cDNA. Amplifications were conducted using an Applied Biosystems 7300 Real Time PCR system and sequence detection software (version 1.4). All samples for qPCR were run in triplicate. The relative quantitation software package was used to determine fold changes in expression for *DEFB4B* and *S100A9*. The software uses the comparative C_T_ (ΔΔ C_T_) method. All fold changes reported here are based on *RACK1* as the reference gene for normalization, and are relative to expression of the target gene in a single non-laminitic control case (sample 1; Table 1). The qPCR analyses were limited to *DEFB4B* and *S100A9* because the other target genes were expressed at undetectable levels in the non-laminitic controls. Two primers were used to confirm the expression of *DEFB4B* (Table 3).

The Wilcoxon-Mann-Whitney rank sum test was used to compare gene expression fold changes between non-laminitic controls and individual disease stages. The analyses were performed using Kaleidagraph software (version 4.5.2).

### In situ hybridization

The entire 345 bp mRNA sequence of *DEFB4B* (NM_001081887.1) was used to synthesize a corresponding cDNA sequence. To facilitate cloning, 5’ NotI and 3’ XhoI restriction sites were included in the synthesized DNA fragment. The synthesized DNA was cloned into pBluescript SK (+) vectors at the multi-cloning site, which includes T7 and T3 RNA polymerase promoter sequences. Gene synthesis, cloning, and verification of sequence were performed by Genscript (genscript.com). Digoxigenin (DIG)-labeled riboprobes were synthesized from T7 (for antisense probe synthesis) and T3 (for sense probe synthesis) promoters using MEGAscript Transcription kits (Ambion, Thermo Fisher Scientific) according to the manufacturer’s protocol.

In situ hybridization was performed using standard methods [62] as described previously [7], modified to include antibody incubation at room temperature. Briefly, tissue sections were deparaffinized in xylene, followed by rehydration in a graded ethanol series (100%, 75%, 50% and 25%) and digested for 5 min with proteinase K (10μg/mL; Ambion). Tissue sections were allowed to hybridize overnight in a humid chamber at 65°C with 1 ng/µL of sense (negative probe) or antisense (positive probe) DIG-labeled riboprobes in hybridization buffer containing 50% formamide. After washes in saline-sodium citrate buffer, the sections were incubated with alkaline phosphatase-conjugated anti-DIG Fab fragments (#11093274910, 1:3000, SigmaAldrich, St. Louis, MO, USA) in a humid chamber for 1 h at 37°C. After washing in PTB (Phosphate Buffered Saline + 0.2% Triton x-100 + 0.1% BSA), labeled probe was visualized using NBT/BCIP substrate (Roche Diagnostics, Indianapolis, IN, USA) resulting in a blue/purple precipitate. PTw buffer (Phosphate Buffered Saline + 0.1% Tween) was used to stop the reaction. Sections were mounted in 80% glycerol/PTw.

Sections from PASH or in situ hybridization were imaged using a Nikon Nti microscope, a Nikon DS /Ti2 color camera and Nikon Elements software (Nikon Instruments, Inc., Melville, NY, USA). Images were collected using 4X or 10X objectives. Images collected at 4X were manually stitched together using Adobe Illustrator, allowing visualization of entire sections.

#### Immunofluorescence

Tissue sections were also used to localize calprotectin using mouse antibody ab22506 (1:5000; Abcam) recognizing S100A9, followed by goat anti mouse IgG-Alexa 488 (1:100), based on a protocol described previously [59]. The calprotectin antibody has been used previously in equine tissues [16; 52; 63]. Rhodamine-tagged wheat germ agglutinin (12.5 µg/ml; Vector Laboratories) was used as a counterstain to outline cell membranes and extracellular matrix, facilitating identification of lamellar microanatomy [58]. Sections were imaged using a Zeiss 880 confocal microscope system including a Zeiss Axio Observer inverted microscope stand and 25X PlanApo objective (0.8 NA), controlled by Zen software (version 2.1). Large areas of tissue were imaged using the tile scan feature, typically set to 0% overlap between sections. The tiled sections were stitched together using Zen software (version 2.1).

## Results

### IL-17 Receptor Subunit A Expression

Before examining downstream effectors of the IL-17 pathway, we first determined that *IL-17RA*, encoding one half of the IL-17 receptor heterodimer, was detectable in most of the samples examined (Fig 3). The receptor subunit was detected in both non-laminitic and laminitic samples.

**Fig 3.**
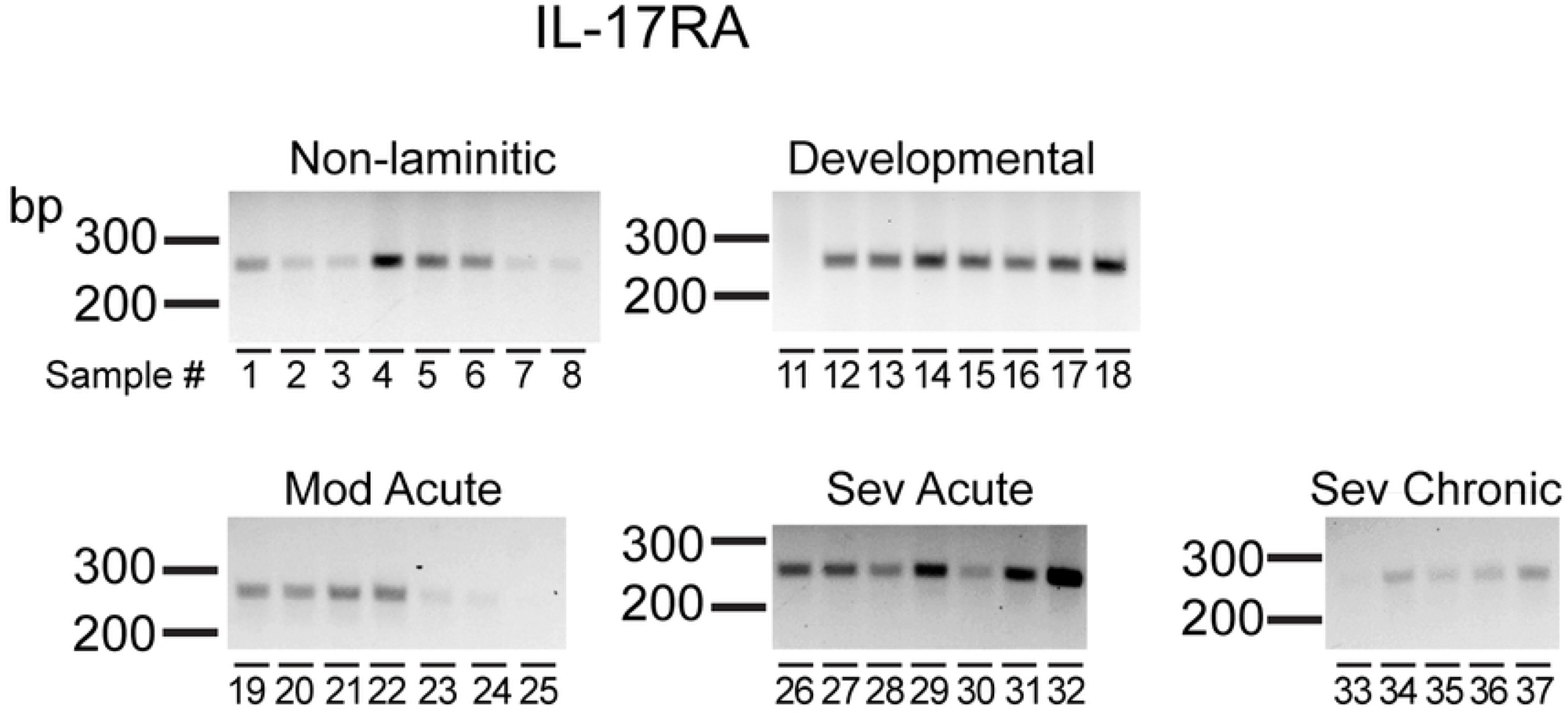
IL-17RA is expressed in equine lamellar tissue. RT-PCR products amplified from equine lamellar tissue isolated from non-laminitic controls and the 4 SLL disease states are shown as indicated. Base pair lengths are indicated to the left of gels. Most samples expressed detectable levels of IL-17RA. Samples numbers are listed in Table 1.

### DEFB4B Expression

Given the high expression of ß defensin 4 (human gene name *DEFB4*) in IL-17-treated human keratinocytes [34], we used qPCR to examine relative expression levels of *DEFB4B* mRNA in SLL samples compared to non-laminitic controls (Fig 4). Note that the equine *DEFB4B* sequence was formerly identified as ß defensin 1 [61]. As shown in Fig 4A, *DEFB4B* expression was upregulated at all disease stages compared to the non-laminitic controls. Modest upregulation was detected in samples from developmental and moderate acute stages. More robust upregulation was present in either severe acute or severe chronic samples. These results were confirmed using a second primer pair (Table 3) for both developmental and severe acute samples (Fig 4B). End points from the qPCR experiment (primer pair 1) are shown in Fig 4C, confirming that primers amplified a sequence of the expected size (Table 3).

**Fig 4.**
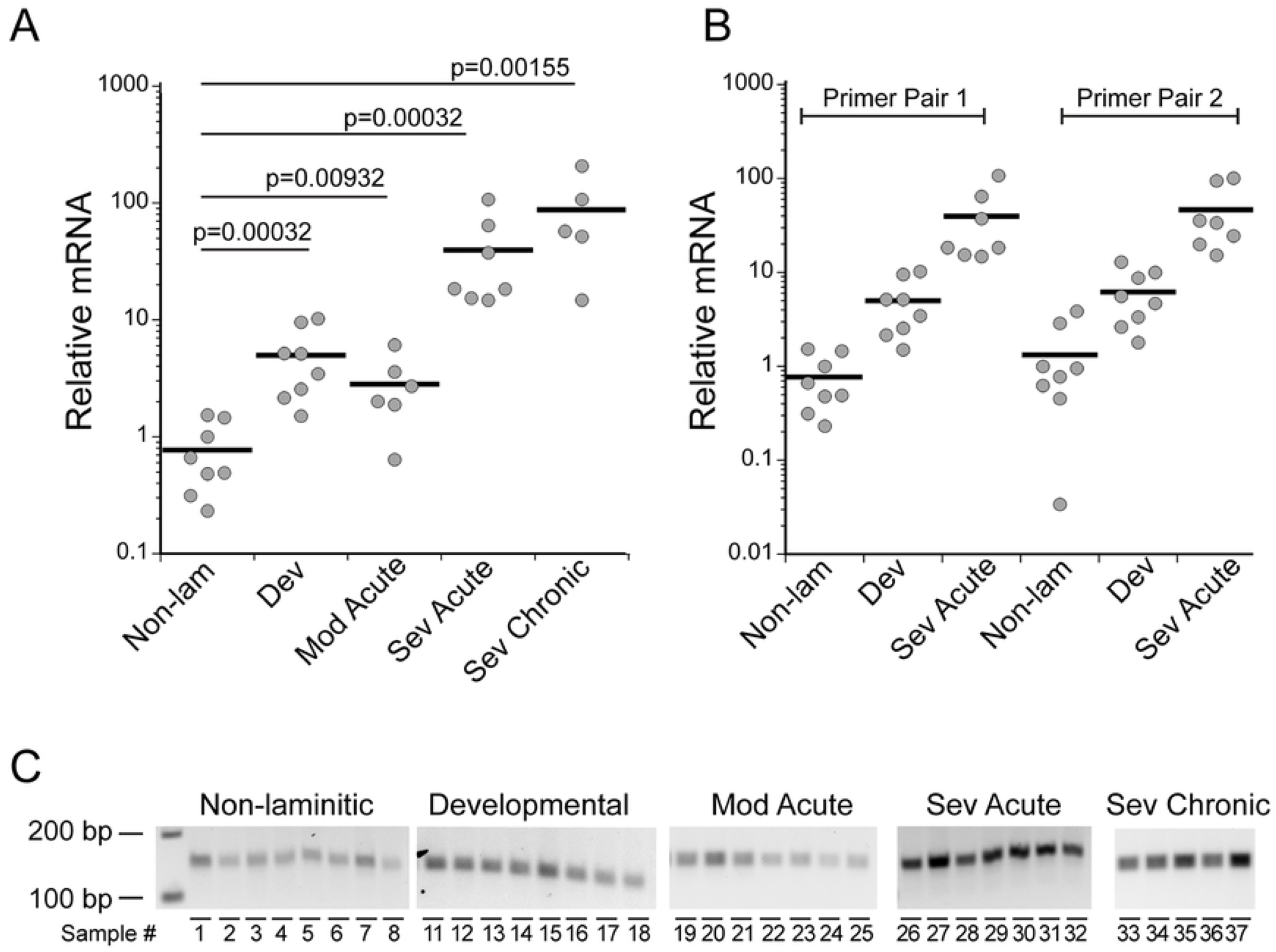
Equine *DEFB4B* expression was upregulated in SLL. (A,B) Relative *DEFB4B* gene expression measured by qPCR (see Methods) in SLL disease states (developmental (Dev), moderate acute (Mod Acute), severe acute (Sev Acute), and severe chronic (Sev Chronic)) compared to non-laminitic controls (Non Lam) as indicated. (A) Relative mRNA levels are shown from qPCR using primer pair 1. (B) Upregulation of *DEFB4B* was verified with primer pair 2. Primer sequences are listed in Table 3. The p values are derived from Wilcoxon-Mann-Whitney rank sum tests. (C) End points from qPCR show amplimers of the predicted 149 bp for primer pair 1. Base pair lengths are indicated to the left of gels and sample numbers are indicated below each lane. Sample information is listed in Table 1.

### Expression of other Effectors of IL-17-dependent signals

As shown in Fig 5, *S100A9, MMP9, S100A8*, and *PTGS2* (common name, COX2, see Table 3) were expressed in severe acute SLL samples, while nearly undetected in the non-laminitic samples. For *S100A9*, the severe acute SLL samples showed significant upregulation of gene expression by qPCR (Fig 5B). The low or undetected expression in the non-laminitic samples for *MMP9, S100A8* or *PTGS2* made it impossible to calculate a fold change for these genes, but they were all clearly expressed in most, or all, of the severe acute samples examined. The reference gene, *RACK1*, is also shown in Fig 5A. All amplimers were detected at the correct size (Table 3).

**Fig 5.**
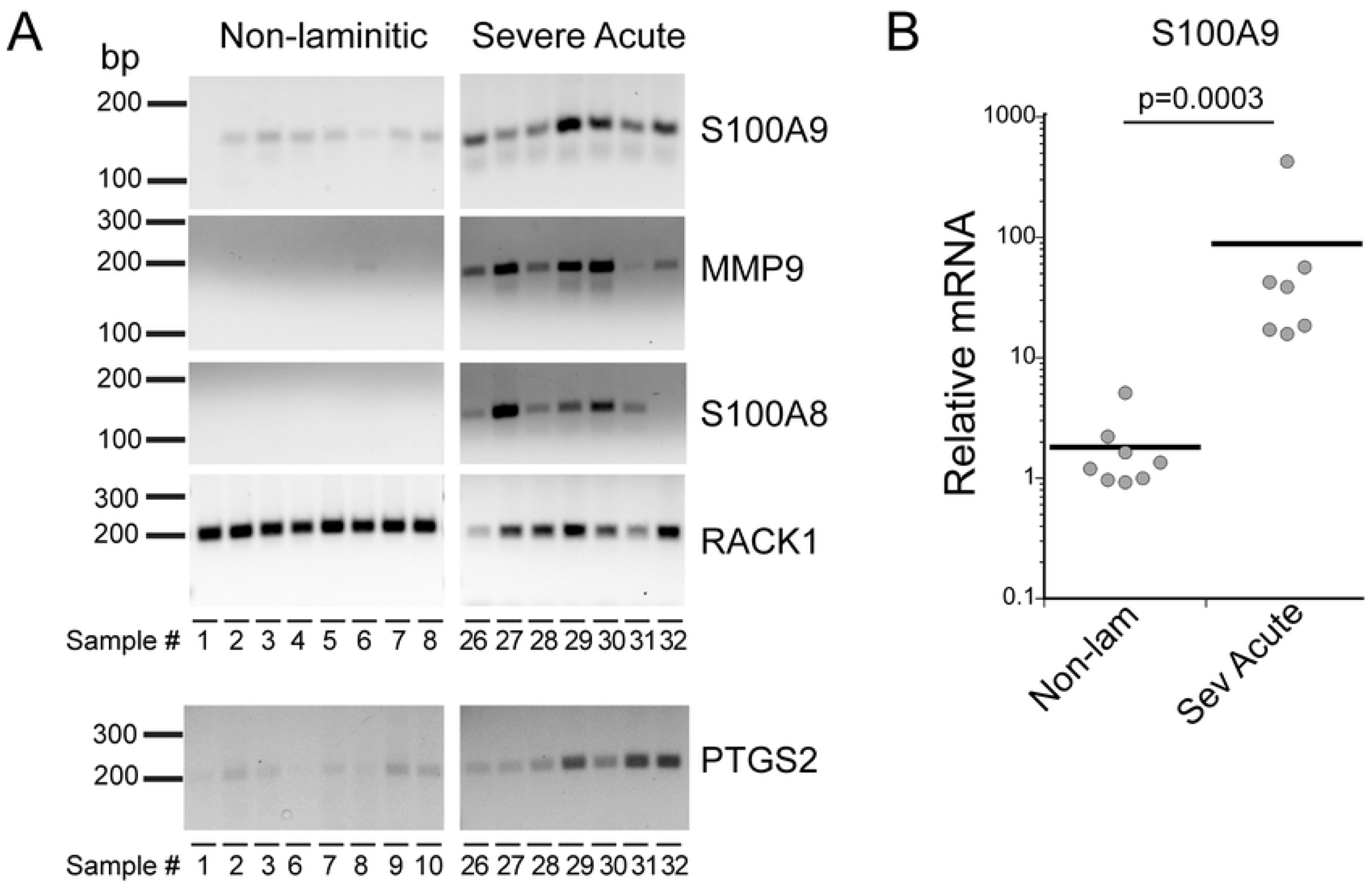
Four IL-17 target genes were expressed in most cases of severe acute SLL. (A) Amplimers are from either the qPCR end points (*S100A9, RACK1*) or from RT-PCR amplification, using the primer sequences listed in Table 3. *RACK1* serves as the reference gene for qPCR. Base pair lengths are indicated to the left of gels and sample numbers are indicated below each lane. Sample information is listed in Table 1. (B) *S100A9* was significantly upregulated in severe acute SLL. The p value was calculated by Wilcoxon-Mann-Whitney rank sum test.

Expression of additional IL-17 target genes in severe acute SLL are shown in Fig 6A for *CCL2, CxCL8, TNFα, IL6* and *MMP1*. In the non-laminitic controls, gene expression was undetected, or very weakly detected in all cases. For the severe acute cases, gene expression was detected in some, but not all cases examined. An additional chemokine, *CCL20*, was weakly detected in non-laminitic controls and not detected in severe acute disease samples (Fig 6B).

**Fig 6.**
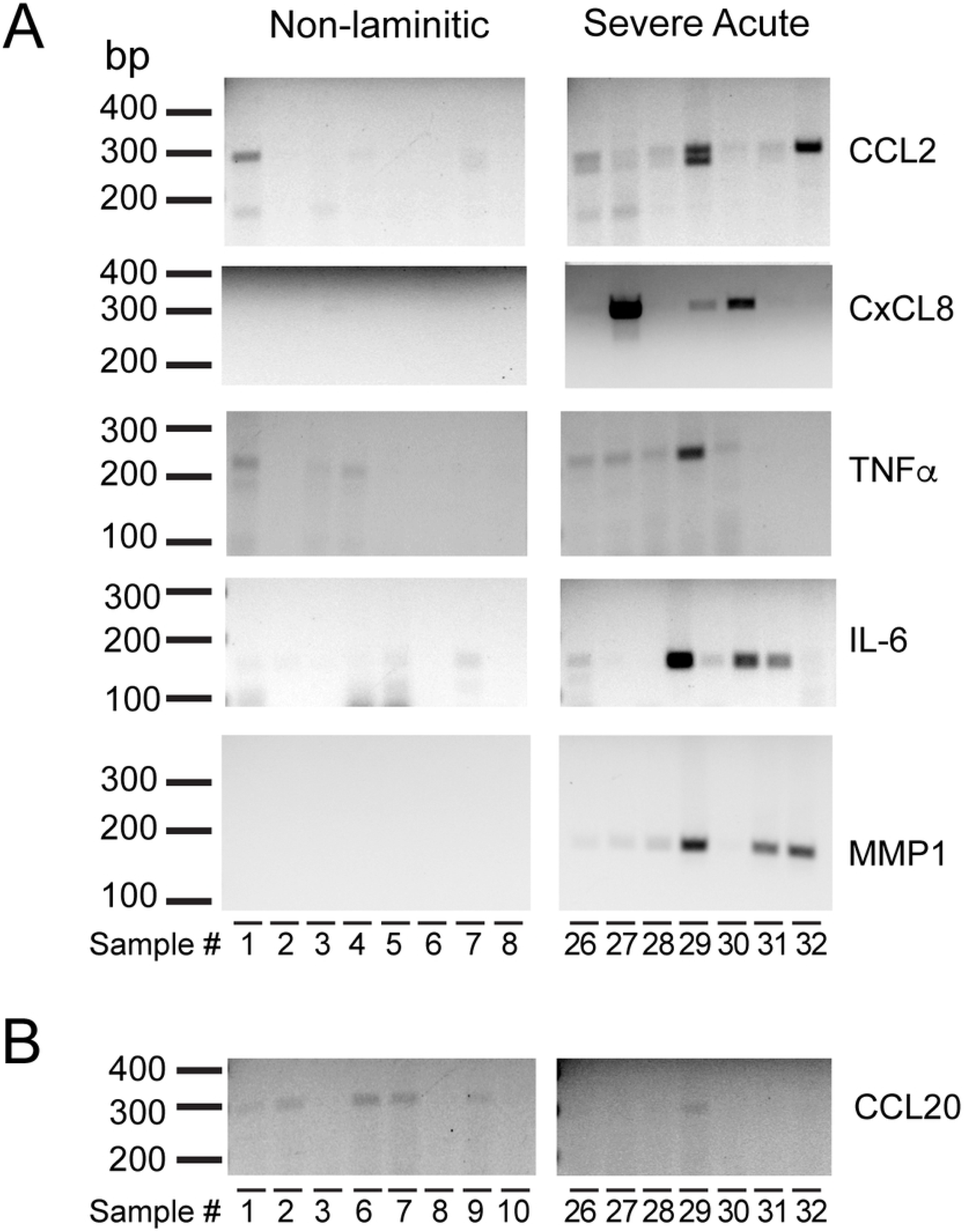
The expression of five IL-17 target genes was detected in some, but not all, severe acute SLL cases. (A) Amplimers from RT-PCR for *CCL2, CxCL8, TNFα, IL6*, and *MMP1* are shown for both non-laminitic controls and severe acute SLL. (B) *CCL20* was not detected in severe acute SLL. Base pair lengths are indicated to the left of gels and sample numbers are indicated below each lane. Sample information is listed in Table 1.

Gene expression was also examined in samples from the other disease stages. Several of the IL-17 target genes were also detected in some, but not all, samples from developmental and moderate acute disease stages (S1 Fig). Most of the developmental samples expressed *PTGS2* and *IL6*, with fewer samples expressing detectable amplimer bands for *CCL2* or *TNFα*. The moderate acute samples showed detectable expression of *PTGS2, CCL2, TNFα*, and *IL6*. For the severe chronic cases, 4 IL-17 target genes, *MMP9, S100A8, CCL2* and *MMP1*, were examined and these were expressed in the majority of severe chronic cases (S1 Fig).

### Localization of cells expressing DEFB4B and calprotectin in SLL

To address whether IL-17 target genes are expressed by keratinocytes of the epidermal lamellae, we examined expression of two IL-17 target genes. *DEFB4B* expression was localized by in situ hybridization to detect the mRNA, and calprotectin was localized by immunofluorescence using a commercial antibody recognizing S100A9. Sequential sections were used for the two localizations and compared to PASH stained sections. Fig 7 shows tissue sections from a severe acute case stained by PASH (Fig 7A), and corresponding sections to localize *DEFB4B* expression (Fig 7B) and calprotectin (Fig 7C). The two boxed regions are shown at higher magnification in Fig 8. For both regions, *DEFB4B*- and calprotectin-expressing cells colocalize to the same areas, primarily suprabasal keratinocytes of the epidermal lamellae. These areas also include cells that aberrantly stain positively with PAS (arrows in Fig 8). Additional images from a second severe acute case and a severe chronic case are shown in S2 Fig; these also demonstrate co-localization of PAS-positive cells, *DEFB4B* and calprotectin to the same areas of the tissue, and are detected primarily in suprabasal keratinocytes. The location of *DEFB4B* and/or calprotectin positive keratinocytes varied between individual cases and ranged from cells found in abaxial regions (S2 Fig), axial regions (Fig 7), or in regions adjacent to gross tissue damage and loss of tissue integrity (Figs 7 and 8).

**Fig 7.**
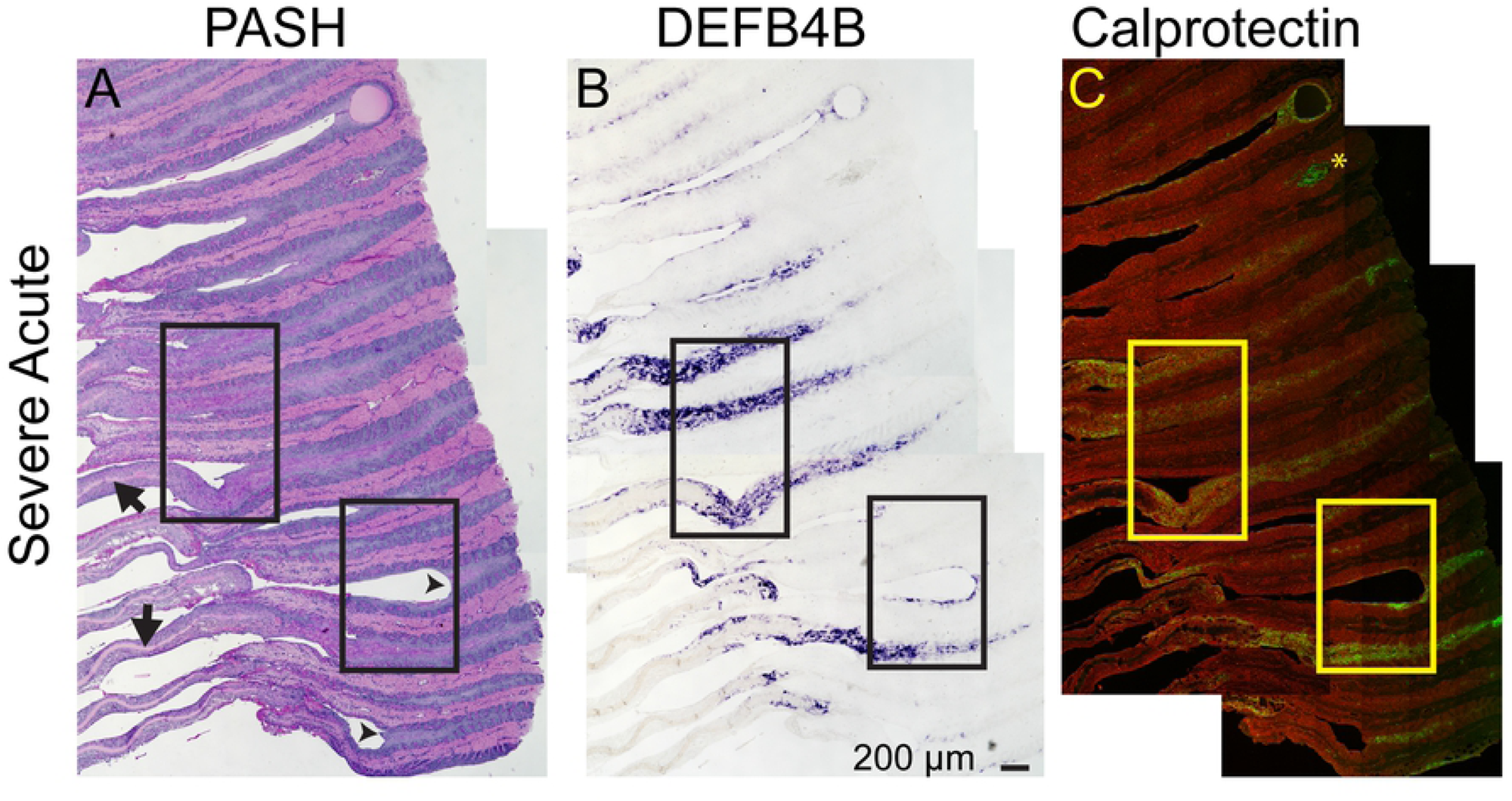
*BDEF4B* and calprotectin were expressed by lamellar keratinocytes in severe acute SLL. Images shown are from sequential FFPE lamellar tissue sections labeled by PASH staining, *DEFB4B* in situ hybridization to localize mRNA (purple color), and calprotectin (antibody recognizing S100A9 protein; green color) by immunofluorescence (n=3 using samples 27, 32, and 38; images from sample 32 shown, see S2 Fig for images from sample 27 (severe acute) and 38 (severe chronic). Arrows in the PASH image mark locations of the centralized keratinized axis (KA) of the PEL, which have been displaced or “stripped” axially, from the axial remnants of the epidermal lamellae (arrowheads). Asterisk in the calprotectin image marks red blood cell autofluorescence. The calprotectin image also includes rhodamine-tagged wheat germ agglutinin (red) as a counterstain to mark both extracellular matrix and cell membranes. Each panel is a composite of multiple images tiled together (see Methods). The two boxed regions are shown at higher magnification in Fig 8. Sections are oriented with the abaxial region to the left and the axial region to the right.

**Fig 8.**
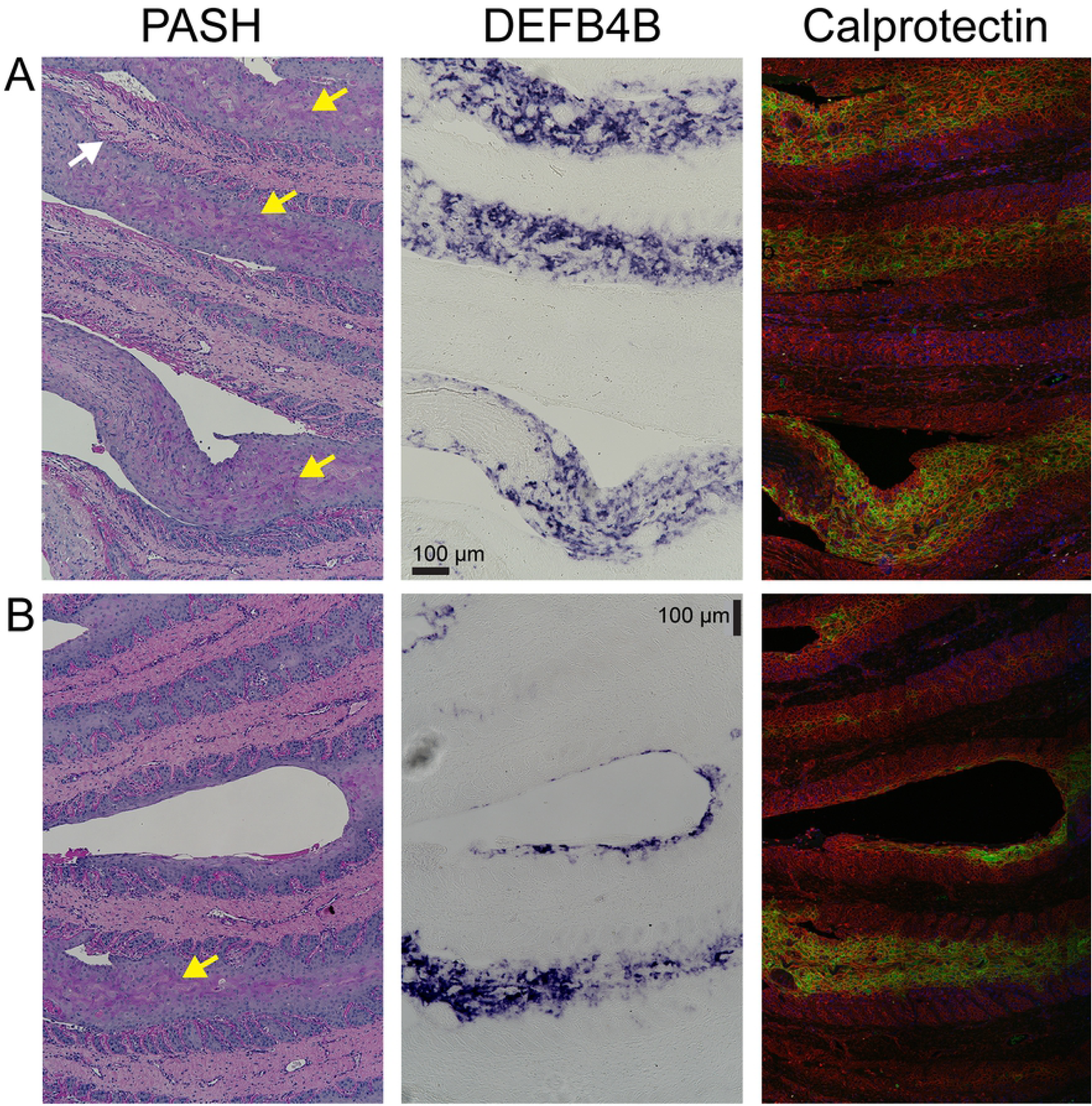
*DEFB4B* and calprotectin localize to suprabasal keratinocytes in areas that also stain aberrantly positively with PAS. Regions boxed in Fig 7 are shown at higher magnification for a severe acute SLL sample. *DEFB4B* mRNA was localized by in situ hybridization (purple color product) and calprotein protein by immunofluorescence (green). The calprotectin image also includes rhodamine-tagged wheat germ agglutinin (red) as a counterstain to mark both extracellular matrix and cell membranes. PAS positive (purple) regions are marked by yellow arrows. PAS also highlights the basement membrane separating dermal and epidermal layers (white arrow). Panel A corresponds to the upper box in Fig 7, Panel B corresponds to the lower box.

Several controls confirmed the specificity of localizations for *DEFB4B* mRNA or calprotectin. First, neither *DEFB4B* nor calprotectin were detected in sections from non-laminitic tissues (S3 Fig). Second, a sense probe generated from the *DEFB4B* sequence did not bind tissue sections other than a weak binding to some cell nuclei (S4 Fig). Suprabasal cells within normal epidermal lamellae do not stain positively with PAS and have extremely limited stratification (S3 Fig).

## Discussion

We found significant evidence for upregulation of the IL-17 pathway in lamellar tissue isolated from horses euthanized for SLL based on the detection and upregulation of 10 out of 11 IL-17 target genes examined (Figs 4 - 6). Additionally, expression of the IL-17R subunit A was detected in both non-laminitic controls and in SLL lamellar tissue (Fig 3), consistent with findings in human keratinocytes that the IL-17 receptor is constitutively expressed [31]. For most IL-17 target genes, expression was undetectable in the non-laminitic samples, and therefore it was not feasible to calculate a gene expression fold change in the SLL samples. Two genes, *DEFB4B* and *S100A9*, were detected at sufficient levels in the non-laminitic tissue for relative quantitation. Both genes were expressed at ∼ 20-100 fold higher levels in tissue from the severe acute stage of disease, compared to that in the non-laminitic controls. Although several markers, including *DEFB4B, S100A9, S100A8, PTGS2* and *MMP9*, were expressed in most or all severe acute samples, additional IL-17 target genes were detected in some, but not all of the severe acute cases examined (Figs 5 and 6). It is possible that the differences in expression of IL-17 target genes among the severe acute samples reflects variation in a specific gene’s expression over time as the disease progresses, or as individual cells respond and adapt to receptor activation. There could also be an underlying genetic variability among horses, as suggested for “non-responders” to laminitis induction in experimental models [46; 48]. Additionally, we failed to detect significant expression of *CCL20* (Fig 6), perhaps indicating a difference in an IL-17 pathway response due to species or tissue differences. Overall, the detectable expression of 10 out of 11 downstream target genes in severe acute SLL tissue supports the hypothesis that lamellar tissue is responding via a similar IL-17 inflammatory pathway to that identified in human and animal models of auto-inflammatory diseases of the skin and extracutaneous sites, including psoriasis/psoriatic enthesis-arthritis and skin cancer, as well as the wound healing response in human or mouse skin [18; 19; 21; 28].

In horses, involvement of the IL-17 pathway has also been identified in the syndrome of recurrent airway obstruction (RAO), a chronic pulmonary inflammatory condition proposed as a naturally-occurring animal disease model for human occupational asthma due to similarities in clinical and pathological features [64; 65]. Both diseases appear to involve an excessive innate immune response to environmental particulate exposure [66]. Bronchoalveolar lavage fluid cells from horses with chronic RAO show increased gene expression of IL-17, but the immune and epithelial cell types involved have not been characterized [67-69]. Korn et al. [64] detected differential gene and protein expression in the mediastinal (pulmonary-draining) lymph nodes from RAO-affected horses that was consistent with an IL-17 response. These studies support a role for the IL-17 pathway in the equine innate immune response and inflammatory disease pathogenesis.

Whether the observed activation of the IL-17 pathway in SLL is part of a mechanism driving disease progression or a downstream response to tissue damage is presently unknown. Supporting the idea that IL-17-dependent inflammation contributes to disease is the observation that several IL-17 target genes examined here were upregulated at the developmental or moderate acute stages. The most prominent gene was *DEFB4B*, which showed an ∼2 - 10 fold increased expression level at developmental and moderate acute stages compared to non-laminitic controls (Fig 4). *PTGS2* and *IL6* were also clearly expressed at the developmental stage (S1 Fig). These data support the hypothesis that the IL-17 pathway is upregulated early in disease progression. Alternatively, IL-17-dependent inflammation may occur as part of a response to cell stress and cell death, which together will reduce epidermal function and contribute to loss of mechanical support. The murine model of EAE supports this idea, where IL- 17 pathway activation is induced by hypoxia and perpetuated through HIF-1*α* activation [53]. For equine laminitis, the experimental model to mimic SLL showed induction of HIF-1*α* during very early stages of disease, before horses showed clinical signs or histological lesions of laminitis, consistent with the idea that metabolic stress and hypoxia may contribute to equine SLL [48]. Although inflammatory markers typically increased in other models of laminitis were not increased in their study, it would be interesting to investigate whether there was evidence of IL-17 pathway activation, an inflammatory pathway that was not fully evaluated. Conversely, it would be beneficial to evaluate HIF-1*α* induction in the cohort of spontaneously occurring cases of SLL reported here, representing various stages of disease development and severity. If results from the two studies corroborate, one could hypothesize that unequal loading of the limbs may predispose the lamellae to tissue hypoxia, HIF-1*α* activation and subsequent upregulation of the IL-17 pathway. Although initially the induction of these pathways could mimic wound healing in skin, where the inflammatory process functions after the damage has occurred, further lamellar damage and IL-17 inflammation may form a feed forward loop, driving exacerbation of hypoxia, tissue dysplasia and irreversible progressive loss of tissue function.

The IL-17 pathway upregulation observed here in natural cases of SLL is also likely present in laminitis associated with other risk factors. In models of SRL, one or more target genes of the IL-17 pathway have been identified as genes upregulated after experimental induction of laminitis [5; 15; 38; 39; 42-46; 52; 70]. One study in an SRL experimental model also hints that the IL-17 pathway may contribute to disease onset or progression. In this experimental system, laminitis development was blocked by continuous digital hypothermia; this treatment also blocked upregulation of IL6 and PTGS2, two targets downstream of IL-17 activation [43]. In the hyperinsulinemia/euglycemic clamp model of EL, the cohort of genes examined included few downstream targets of IL-17 activation, but these studies identified both *MMP9* [40] and *TNFα* [41] upregulation. Several studies of naturally occurring cases of laminitis have described increased *MMP9* but the initiating trigger(s) in these natural cases was not described [51; 70]. Taken together, data from models of SRL and EL, as well as the results presented here for SLL, indicate that IL-17 based inflammation is likely a shared feature among the three major triggers of laminitis, although this conclusion needs to be confirmed experimentally by a more comprehensive investigation of IL-17 downstream markers.

Activation of the IL-17 pathway is likely to be relevant to the loss of cell adhesion, cell stretch, and altered epidermal differentiation and osteolysis of the distal phalanx that are key features of laminitis pathogenesis [11; 12; 13; 57; 71]. Changes in keratinocyte differentiation in laminitis are evidenced by both histologic lesions, such as epidermal hyperplasia, metaplasia, orthokeratosis, lamellar basal cell polymorphism, and SEL morphologic lesions [9; 10-14; 40; 59; 71; 72] and in molecular changes that impact the epithelial phenotype [57; 73]. The morphologic changes in the epidermal lamellae are suggestive of changes in the expression of cytoskeletal and cell adhesion molecules. We are currently investigating the impact of laminitis and IL-17 signaling on keratin isoform expression since this is an important feature of human psoriasis and the associated epidermal hyperplasia [74-76] as well as the human nail disease, pachyonychia congenita, characterized by hyperkeratosis of the nail bed [77]. The increase in expression of *DEFB4B* in laminitic tissue reported here could contribute to a loss of epidermal lamellar cell adhesion and the epithelial phenotype in favor of a more proliferative and migratory phenotype, as is the case for human keratinocytes exposed to β-defensins [78]. The ability of β- defensins to promote cell migration and proliferation contributes to wound healing, but could also contribute to laminitis pathogenesis through loss of cell adhesion and epithelial integrity. Suppression of the canonical Wnt signaling pathway has been associated with decreased expression of β catenin and integrin β4 and loss of cell-cell and cell-matrix adhesion in the lamellar basal cells from a SRL model [73]. Interestingly, IL-17-mediated suppression of Wnt signaling is also implicated in the promotion of destructive osteolysis and aberrant osteogenesis that are characteristic of inflammatory osteoarthritis and IL-17 has direct osteoclastogenic effects [75; 76]. We previously demonstrated that osteolysis of the distal phalanx is present in natural cases of acute and chronic laminitis with various associated risk factors [11]. It is therefore possible that IL-17 and Wnt signaling pathways are important in the pathogenesis of both the epidermal and bone lesions of laminitis with the IL-17 produced in lamellar tissue driving bone loss in the adjacent distal phalanx, as occurs in human natural cases and murine models of psoriasis [76; 77].

In human psoriasis, keratinocytes of the skin express the IL-17 receptor, respond to IL- 17 by expression of multiple IL-17 target genes, and are thought to make a major contribution to disease progression [18; 31; 34; 37]. We found that keratinocytes of the equine lamellae express both *DEFB4B* and calprotectin (S100A8/S100A9 dimer), as shown in Figs 7 and 8. Several studies using experimental models of SRL also support a role of lamellar keratinocytes in producing IL-17 target genes. First, calprotectin was also localized to keratinocytes in a model of SRL [47]. Second, epithelial cells isolated by laser microdissection showed upregulation of several IL-17 pathway target genes measured by RNA-seq [15]. Additionally, cultured equine skin keratinocytes exposed to a combination of lipopolysaccharide and hypoxia significantly upregulated expression of *PTGS2* [45]. Although other cell types in the equine lamellae or DP may also express IL-17 target genes, keratinocytes are likely the major cell type expressing IL- 17 target genes in SLL, and possibly laminitis more generally, given their abundance within the foot. The epidermal surface area of the lamellar epithelium of each hoof capsule is approximately 0.8 m2 [78], compared to about 1.5 2 m2 for the total body surface area of human skin [79]. Local signals within each foot of the horse have the potential to reach keratinocytes at a number approaching that present within the entire skin of a human.

We found that *DEFB4B* and calprotectin were present in some, but not all, suprabasal keratinocytes. Additionally, the response varied regionally, with some PEL or SEL regions showing high expression compared to nearby PELs or SELs (Fig 7). From localizations in sequential sections, cells in the same areas expressed both *DEFB4B* and calprotectin (Figs 7 and 8). Cells in these regions also stained positively by PAS, a histological stain that binds glycoproteins or glycogen (Figs 7 and 8). PAS-positive lamellar epithelial cells are not present in healthy adult lamellae [11] (S3 Fig), but are observed in laminitic lamellae in regions that demonstrate acanthosis (areas with expansion of the suprabasal cell layers), resembling stratified squamous tissue with development of discernable desmosomal attachments (see also [11]). Whether PAS-positive staining is always correlated with *DEFB4B* or calprotectin expression has not been examined, but the co-localizations indicate that the same lamellar epithelial cell region is also responding to IL-17 receptor activation. It is notable that these suprabasilar cells are in the region where the PEL keratinized axis and some SEL suprabasal cells have detached from the SEL basal and remaining suprabasal cells, resulting in detachment of the hoof capsule from the DP.

Studies detecting cell surface markers are required to determine which cell types are producing IL-17 and what is triggering IL-17 release within the lamellar environment to activate IL-17 receptors and expression of downstream targets. Given that IL-17-dependent signals result in a feed forward loop that is able to self-amplify locally within tissues [20; 23; 28; 37], it will be challenging to sort out an order of events in a complex tissue that is typically only accessible after euthanasia. Additionally, several signals produced by IL-17-activated cells, including TNF*α* and IL-1ß, can act synergistically with IL-17 to increase the response, and these signals can also be produced in response to other cytokine signals in addition to IL-17 [21; 28; 34]. The IL-17-dependent pathway also may be linked to mTORC1 activation, suggesting additional feed-forward signals. Active mTORC1 has been demonstrated in models of either EL or SRL [80; 81]. Importantly, digital hypothermia blocks mTORC1 activation and laminitis development in these experiments [80; 81]. IL-17 dependent signals are potential upstream activators of mTORC1, where IL-6, which can be produced by multiple cell types in response to signals including IL-17 stimulation of keratinocytes [80], relays signals to activate mTORC1 [80]. Active mTORC1 can also stimulate IL-17 expression [82], consistent with a feed-forward amplification loop.

Assuming IL-17 pathway activation contributes to disease progression in equine laminitis, as it does in human skin diseases, blocking the pathway at upstream points could be advantageous, since it could shut down the feed-forward loop. Inhibition of either mTORC1 or IL-17 receptor activation, or both simultaneously, should inhibit signal amplifications through both pathways. Inhibiting IL-17 signaling in the horse will not be as simple as applying human therapies since the most successful psoriasis treatments use monoclonal antibodies to block IL- 17 receptor activation [21]; these antibodies may not bind the equine IL-17 receptor, may be antigenic in equines, and are likely cost-prohibitive. Local inhibition of IL-17 and/or mTORC1 would be desirable, and could be possible using an adeno-associated viral vector to deliver genes for inhibitory cytokines to reduce inflammation, as demonstrated recently [83], or other genes able to inhibit IL-17 and/or mTORC1 activation.

## Acknowledgements

We gratefully acknowledge the following colleagues: Caitlin Armstrong isolated the RNA used in this study; Karie Durynski cut histology sections; Dr. Jamie Havrilak and Dylan Faltine-Gonzalez shared reagents for in situ hybridization and graciously provided technical advice throughout this project; Dr. Michael Layden shared his Nikon microscope and thermocyclers.

## Supporting Information

**S1 Fig. Expression of 8 IL-17 target genes in developmental and moderate acute SLL disease stages and of 4 target genes in severe chronic SLL detected by RT-PCR**. (A) *PTGS2, CCL2, TNFα* and *IL6* were expressed in some of the developmental and moderate acute samples examined. *CCL20* was not detected at these disease stages. (B) For the severe chronic cohort of cases, expression was examined for four IL-17 target genes. *MMP9, S100A8, CCL2* and *MMP1* were detected in most severe chronic samples. Base pair lengths are indicated to the left of gels and sample numbers are indicated below each lane. *DEFB4B* expression at all disease stages is shown in Fig 4. Sample information is listed in Table 1.

**S2 Fig. Two additional SLL cases showing regional co-localizations of PAS positive cells, DEFB4B (purple product; in situ hybridization) and calprotectin (green; immunofluorescence)**. A. Severe acute case (Sample # 27). Boxed region for PASH and DEFB4B panels are shown below for calprotectin. Three PELs are marked in both the DEFB4B and calprotectin images. B. Severe chronic case (Sample # 38). Boxed region is shown in the bottom row and includes calprotectin localization (green). Sequential FFPE lamellar tissue sections were used for each label. The calprotectin images include rhodamine-tagged wheat germ agglutinin (red) as a counterstain to highlight extracellular matrix and cell membranes.

**S3 Fig. Non-laminitic control tissues do not express *DEFB4B* or calprotectin**. Sequential FFPE lamellar tissue sections from sample #3 stained for PASH (left), in situ hybridization to localize *DEFB4B* expressing cells (middle) or calprotectin (green; right) (n=3 using samples 1, 3 and 5). *DEFB4B* expression was not detected in the non-laminitic sections as evidenced by the lack of purple color product. Non-laminitic tissue sections do not show detectable calprotectin expression (green). Extracellular matrix and cell membranes are marked in red by the counterstain, rhodamine-tagged wheat germ agglutinin (WGA). The darker shadows in the rhodamine signal that appear as a window pane-like pattern are an artifact produced by stitching a series of images together (Methods). Images are oriented with the abaxial region is toward the top and the axial region toward the bottom. A higher magnification PASH image is shown to highlight PAS-positive basement membranes (arrow) and lack of PAS positive cells within SELs (asterisk).

**S4 Fig. Sense control probe does not hybridize to lamellar tissue**. An additional section from the tissue shown in Figs 7 and 8 is shown after hybridization with a sense probe as a control for in situ hybridization. A low magnification composite for the anti-sense probe is shown to the left for comparison (this composite is also shown in Fig 7). Hybridization with the sense probe shows only dim purple reaction product. The boxed regions are shown at higher magnification after a 90° clockwise rotation. The small purple dots mark some cell nuclei, but this pattern is not observed with the anti-sense probe.

## References

[1] Pollitt CC. The anatomy and physiology of the suspensory apparatus of the distal phalanx. Vet Clin Equine. 2010;26(1): 29–49.

[2] Pollitt, CC and Collins SN. The suspensory apparatus of the distal phalanx in normal horses. Eq Vet J. 2016;48(4): 496–501.

[3] Bragulla H, Hirschberg RM. 2003. Horse hooves and bird feathers; Two model systems for studying the structure and development of highly adapted integumentary accessory organs - The role of the dermo-epidermal interface for the micro-architecture of complex epidermal structures. J Exp Zool B Mol Dev Evol. 2003;298(1): 140–51.

[4] Leise B. The role of neutrophils in equine laminitis. Cell Tissue Res. 2018;371(3): 541–50.

[5] Carter RA, Shekk V, de Laat MA, Pollitt CC, Galantino-Homer HL. 2010. Novel keratins identified by quantitative proteomic analysis as the major cytoskeletal proteins of equine (Equus caballus) hoof lamellar tissue. J Anim Sci. 2010;88(12): 3843–55.

[6] Linardi RL, Megee SO, Mainardi SR, Senoo M, Galantino-Homer. Expression and localization of epithelial stem cell and differentiation markers in equine skin, eye and hoof. Vet Dermatol. 2015;26(4): 213–22.

[7] Armstrong C, Cassimeris L, Da Silva Santos C, Microogullari Y, Wagner B, Babasyan S, et al. The expression of equine keratins K42 and K124 is restricted to the hoof epidermal lamellae of Equus caballus. PLoS One. 2019;14(9): e0219234.

[8] Thomason JJ, McClinchey HL, Faramarzi B, Jofriet JC. Mechanical behavior and quantitative morphology of the equine laminar junction. Anat Rec A Discov Mol Cell Evol Biol. 2005;283(2): 366–79.

[9] Kuwano A, Katayama Y, Kasashima Y, Okada K, Reilly JD. A gross and histopathological study of an ectopic white line development in equine laminitis. J Vet Med Sci. 2002;64(10): 893–900.

[10] Collins SN, Van Eps AW, Kuwano A, Pollitt CC. The lamellar wedge. Vet Clin North Am Equine Pract. 2010;26(1): 179–95.

[11] Engiles JB, Galantino-Homer HL, Boston R, McDonald D, Dishowitz M, Hankenson KD. 2015. Osteopathology in the equine distal phalanx associated with the development and progression of laminitis. Vet Pathol. 2015;52(5): 928–44.

[12] Patterson-Kane JC, Karikoski NP, McGowan CM. Paradigm shifts in understanding equine laminitis. Vet J. 2018;231: 33–40.

[13] van Eps A, Burns TA. Are there shared mechanisms in the pathophysiology of different clinical forms of laminitis and what are the implications for prevention and treatment? Vet Clin North Am Equine Pract. 2019;35(2): 379–98.

[14] Karikoski NP, McGowan CM, Singer ER, Asplin KE, Tulamo R-M, Patterson-Kane JC. Pathology of natural cases of equine endocrinopathic laminitis associated with hyperinsulinemia. Vet Pathol. 2015;52(5): 945–56.

[15] Leise BS, Watts MR, Roy S, Yilmaz AS, Alder H, Belknap JK. Use of laser capture microdissection for the assessment of equine lamellar basal epithelial cell signalling in the early stages of laminitis. Equine Vet J. 2015;47(4): 478–88.

[16] Faleiros RR, Nuovo GJ, Belknap JK. Calprotectin in myeloid and epithelial cells of laminae from horses with black walnut extract induced laminitis. J Vet Intern Med. 2009;23(1): 174–81.

[17] Martin DA, Towne JE, Kricorian G, Klekotka P, Gudjonsson JE, Krueger JG, et al. 2013. The emerging role of IL-17 in the pathogenesis of psoriasis: preclinical and clinical findings. J Invest Dermatol. 2013;133(1): 17–26.

[18] Blauvelt A, Chiricozzi A. The immunological role of IL-17 in psoriasis and psoriatic arthritis pathogenesis. Clinical Rev Allergy Immunol. 2018;55(3): 379–90.

[19] Zhao J, Chen X, Herjan T, Li X. The role of interleukin-17 in tumor development and progression. J Exp Med. 2020;217(1): e20190297. doi: 10.1084/jem.20190297.

[20] Hawkes JE, Yan BY, Chan TC, Krueger JG. Discovery of the IL-23/IL-17 signaling pathway and the treatment of psoriasis. J Immunol. 2018;201(6): 1605–13.

[21] Prinz I, Sandrock I, Mrowietz U. Interleukin-17 cytokines: Effectors and targets in psoriasis - A breakthrough in understanding and treatment. J Exp Med. 2020;217(1): e20191397. doi: 10.1084/jem.20191397.

[22] Rendon A, Schakel K. Psoriasis pathogenesis and treatment. Int J Mol Sci. 2019; 20(6). 1475.

[23] Krueger JG, Brunner PM. Interleukin-17 alters the biology of many cell types involved in the genesis of psoriasis, systemic inflammation and associated comorbidities. Exp Derm. 2018;27(2): 115–23.

[24] Kual S, Singal A, Grover C, Sharma S. Clinical and histological spectrum of nail psoriasis: A cross-sectional study. J Cutan Pathol. 2018l;45(11): 824–30.

[25] Antony AS, Allard A, Rambojun A, Lovell CR, Shaddick G, Robinson G et al. Psoriatic nail dystrophy is associated with erosive disease in the distal interphalangeal joints in psoriatic arthritis: A retrospective cohort study. J Rheumatol. 2019;46(9): 1097–1102.

[26] Xiao M, Wang C, Zhang J, Li Z, Zhao X, Qin Z. IFNgamma promotes papilloma development by up-regulating Th17-associated inflammation. Cancer Res. 2009;69(5): 2010–7.

[27] Wang L, Yi T, Zhang W, Pardoll DM, Yu H. IL-17 enhances tumor development in carcinogen-induced skin cancer. Cancer Res. 2010;70(24): 10112–20.

[28] McGeachy MJ, Cua DJ, Gaffen SL. The IL-17 family of cytokines in health and disease. Immunity. 2019;50(4): 892–906.

[29] Chen X, Cai G, Liu C, Zhao J, Gu C, Wu L, et al. 2019. IL-17R-EGFR axis links wound healing to tumorigenesis in Lrig1+ stem cells. J Exp Med. 2019;216(1): 195–214.

[30] Harper EG, Changsheng G, Rizzo H, Lillis JV, Kurtz SE, Skorcheva I, et al. Th17 cytokines stimulate CCL20 expression in keratinocytes in vitro and in vivo: implications for psoriasis pathogenesis. J Invest Dermatol. 2009;129(9): 2175–83.

[31] Nograles KE, Zaba LC, Guttman-Yassky E, Fuentes-Duculan J, Suarez-Farinas M, Cardinale I, et al. Th17 cytokines interleukin (IL)-17 and IL-22 modulate distinct inflammatory and keratinoctye-response pathways. Br J Dermatol. 2008;159(5): 1092–1102.

[33] Amatya N, Garg AV, Gaffen SL. IL-17 signaling: The yin and the yang. Trends Immunol. 2017;38(5): 310–22.

[34] Chiricozzi A, Guttman-Yassky E, Suarez-Farinas M, Nograles KE, Tian S, Cardinale I, et al. 2011. Integrative responses to IL-17 and TNF-alpha in human keratinocytes account for key inflammatory pathogenic circuits in psoriasis. J Invest Dermatol. 2011;131(3): 677–87.

[35] Jansen PA, Rodijk-Olthuis D, Hollox EJ, Kamsteeg M, Tjabringa GS, de Jongh GJ, et al. Beta-defensin-2 protein is a serum biomarker for disease activity in psoriasis and reaches biologically relevant concentrations in lesional skin. PLoS One. 2009;4(3): e4725.

[36] Kolbinger F, Loesche C, Valentin M-A, Jian X, Cheng Y, Jarvis P, et al. 2017. Beta-defensin 2 is a responsive biomarker of IL-17A-driven skin pathology in patients with psoriasis. J Allergy Clin Immunol. 2017;139(3): 923–32.

[37] Ha H-L, Wang H, Pisitkun P, Kim J-C, Tassi I, Tang W, et al. IL-17 drives psoriatic inflammation via distinct, target cell-specific mechanisms. Proc Natl Acad Sci U S A. 2014;111(33): E3422–31.

[38] Noschka E, Vandenplas ML, Hurley DJ, and Moore JN. Temporal aspects of laminar gene expression during the developmental stages of equine laminitis. Vet Immunol Immunopathol. 2009;129(3-4): 242–53.

[39] Blikslager AT, Yin C, Cochran AM, Wooten JG, Pettigrew A, Belknap JK. Cyclooxygenase expression in the early stages of equine laminitis: a cytologic study. J Vet Intern Med. 2006;20(5): 1191–6.

[40] de Laat MA, Kyaw-Tanner MT, Nourian AR, McGowan CM, Sillence MN, Pollitt CC. The developmental and acute phases of insulin-induced laminitis involve minimal metalloproteinase activity. Vet Immunol Immunopathol. 2011;140(3-4): 275–81.

[41] de Laat MA, Clement CK, McGowan CM, Sillence MN, Pollitt CC, Lacombe VA. Toll-like receptor and pro-inflammatory cytokine expression during prolonged hyperinsulinaemia in horses: Implications for laminitis. Vet Immunol Immunopathol. 2014;157(1-2): 78–86.

[42] Dern K, Watts M, Werle B, van Eps A, Pollitt C, Belknap J. Effect of delayed digital hypothermia on lamellar inflammatory signaling in the oligofructose laminitis model. J Vet Intern Med. 2017;31(2): 575–81.

[43] Dern K, van Eps A, Wittum T, Watts M, Pollitt C, Belknap J. Effect of continuous digital hypothermia on lamellar inflammatory signaling when applied at a clinically-relevant timepoint in the oligofructose laminitis model. J Vet Intern Med. 2018;32(1): 450–8.

[44] Wang L, Pawlak EA, Johnson PJ, Belknap JK, Alfandari D, Black SJ. 2014. Expression and activity of collagenases in the digital laminae of horses with carbohydrate overload-induced acute laminitis. J Vet Intern Med. 2014;28(1): 215–22.

[45] Pawlak EA, Geor RJ, Watts MR, Black SJ, Johnson PJ, Belknap JK. Regulation of hypoxia-inducible factor 1alpha and related genes in equine digital lamellae and in cultured keratinocytes. Equine Vet J. 2014;46(2): 203–9.

[46] Leise BS, Faleiros RR, Watts M, Johnson PJ, Black SJ, Belknap JK. Laminar inflammatory gene expression in the carbohydrate overload model of equine laminitis. Equine Vet J. 2011;43(1): 54–61.

[47] Faleiros RR, Leise BS, Watts M, Johnson PJ, Black SJ, Belknap JK. Laminar chemokine mRNA concentrations in horses with carbohydrate overload-induced laminitis. Vet Immunol Immunopathol. 2011;144(1-2): 45–51.

[48] Gardner AK, van Eps AW, Watts MR, Burns TA, Belknap JK. A novel model to assess lamellar signaling relevant to preferential weight bearing in the horse. Vet J. 2017;221: 62–7.

[49] Watts MR, Hegedus OC, Eades SC, Belknap JK, Burns TA. Association of sustained supraphysiologic hyperinsulinemia and inflammatory signaling within the digital lamellae in light-breed horses. J Vet Intern Med. 2019;33(3): 1483–92.

[50] Black SJ, Lunn DP, Yin C, Hwang M, Lenz SD, Belknap JK. Leukocyte emigration in the early stages of laminitis. Vet Immunol Immunopathol. 2006;109(1-2): 161–6.

[51] Loftus JP, Johnson PJ, Belknap JK, Pettigrew A, Black SJ. Leukocyte-derived and endogenous matrix metalloproteinases in the lamellae of horses with naturally acquired and experimentally induced laminitis. Vet Immunol Immunopathol. 2009;129(3-4): 221–30.

[52] Faleiros RR, Johnson PJ, Nuovo GJ, Messer NT, Black SJ, Belknap JK. Laminar leukocyte accumulation in horses with carbohydrate overload-induced laminitis. J Vet Intern Med. 2011;25(1): 107–15.

[53] Dang EV, Barbi J, Yang HY, Jinasena D, Yu H, Zheng Y et al. Control of T(H)17/T(reg) balance by hypoxia-inducible factor 1. Cell. 2011;146(5): 772–84.

[54] Visser MB, Pollitt CC. Lamellar leukocyte infiltration and involvement of IL-6 during oligofructose-induced equine laminitis development. Vet Immunol Immunopathol. 2011;144(1-2): 120–8.

[55] Godman JD, Burns TA, Kelly CS, Watts MR, Leise BS, Schroeder EL, et al. The effect of hypothermia on influx of leukocytes in the digital lamellae of horses with oligofructose-induced laminitis. 2016;178: 22–8.

[56] Galantino-Homer H, Carter R, Megee S, Engiles J, Orsini J, Pollitt C. The laminitis discovery database. J Equine Vet Sci. 2010;30(2): 101.

[57] Carter RA, Engiles JB, Megee SO, Senoo M, Galantino-Homer HL. 2011. Decreased expression of p63, a regulator of epidermal stem cells, in the chronic laminitic equine hoof. Equine Vet J. 2011;43(5): 543–51.

[58] Clark RK, Galantino-Homer HL. 2014. Wheat germ agglutinin as a counterstain for immunofluorescence studies of equine hoof lamellae. Exp. Dermatol. 2014;23(9): 677–8.

[59] Cassimeris L, Engiles JB, Galantino-Homer H. 2019. Detection of endoplasmic reticulum stress and the unfolded protein response in naturally-occurring endocrinopathic equine laminitis. BMC Vet Res. 2019;15(1): 24.

[60] Wang J. Genome-wide transcriptome analysis of laminar tissue during the early stages of experimentally induced equine laminitis [Doctorate]. College Station, TX: Texas A&M University. 2010.142 p. available from: https://oaktrust.library.tamu.edu/bitstream/handle/1969.1/ETD-TAMU-2010-12-8718/WANG-DISSERTATION.pdf?sequence=2&isAllowed=y

[61] Marth CD, Firestone SM, Glenton LY, Browning GF, Young ND, Krekeler N. Oestrous cycle-dependent equine uterine immune response to induced infectious endometritis. Vet Res. 2016;47(1): 110.

[62] Wolenski FS, Layden MJ, Martindale MQ, Gilmore TD, Finnerty JR. Characterizing the spatiotemporal expression of RNAs and proteins in the starlet sea anemone Nematostella vectensis. Nat Protoc. 2013;8(5): 900–15.

[63] Burns TA, Geor RJ, Mudge MC, McCutcheon LJ, Hinchcliff KW, Belknap JK. Proinflammatory cytokine and chemokine gene expression profiles in subcutaneous and visceral adipose tissue depots of insulin-resistant and insulin-sensitive light breed horses. J Vet Intern Med. 2010;24(4): 932–9.

[64] Korn A, Miller D, Dong L, Buckles EL, Wagner B, Ainsworth DM. Differential gene expression profiles and selected cytokine protein analysis of mediastinal lymph nodes of horses with chronic recurrent airway obstruction (RAO) support an interleukin-17 immune response. PLoS One. 2015;10(11): e0142622.

[65] Poole JA, Romberger MD. Immunological and inflammatory responses to organic dust in agriculture. Curr Opin Allergy Clin Immunol. 2012;12(2): 126–132.

[66] Pirie RS, Dixon PM, Collie DD, McGorum BC. Pulmonary and systemic effects of inhaled endotoxin in control and heaves horses. Eq Vet J. 2001 33(3): 311–8.

[67] Dubrue M, Hamilton E, Joubert P, Lavoie-Kadoch S, Lavoie J-P. Chronic exacerbation of equine heaves is associated with an increased expression of interleukin-17 mRNA in bronchoalveloar lavage. Vet Immunol Immunopathol. 2005;105(1-2): 25–31.

[68] Ainsworth DM, Wagner B, Franchini M, Grunig G, Erb HN, Tan JT. Time-dependent alterations in gene expression of interleukin-8 in the bronchial epithelium of horses with recurrent airway obstruction. Am J Vet Res. 2006;67(4): 669–677.

[69] Padoan E, Ferraresso S, Pegolo S, Castagnaro M, Barnini C, Bargellini L. Real time RT-PCR analysis of inflammatory mediator expression in recurrent airway obstruction-affected horses. Vet Immunol Immunopathol. 2013;156(3-4): 190–199.

[70] Coyne MJ, Cousin H, Loftus JP, Johnson PJ, Belknap JK, Gradil CM, et al. Cloning and expression of ADAM-related metalloproteases in equine laminitis. Vet Immunol Immunopathol. 2009;129(3-4): 231–41.

[71] Pollitt CC. Basement membrane pathology: a feature of acute equine laminitis. Equine Vet J. 1996;28(1): 38–46.

[72] Karikoski NP, Patterson-Kane JC, Singer ER, McFarlane D, McGowan CM. Lamellar pathology in horses with pituitary pars intermedia dysfunction (PPID). Equine Vet J. 2016;48(4): 472–8.

[73] Wang L, Pawlak EA, Johnson PJ, Belknap JK, Eades S, Stalenhoef AF et al. 2013. Impact of laminitis on the canonical Wnt signaling pathway in basal epithelial cells of the equine digital laminae. PLoS One. 2013;8(2): e56025.

[74] Freedberg IM, Tomic-Canic M, Komine M, Blumenberg M. Keratins and the keratinocyte activation cycle. J Invest Dermatol. 2001;116(5): 633–40.

[75] Leigh IM, Navsaria H, Purkis PE, McKay IA, Bowden PE, Riddle PN. Keratins (K16 and K17) as markers of keratinocyte hyperproliferation in psoriasis in vivo and in vitro. Br J Dermatol. 1995;133(4): 501–11.

[76] Lomine M, Freedberg IM, Blumenberg M. Regulation of epidermal expression of keratin 17 in inflammatory skin diseases. J Invest Dermatol. 1996;107(4): 569–75.

[77] McGowan KM, Coulombe PA. Keratin 17 expression in the hard epithelial context of the hair and nail, and its relevance for the pachyonychia congenita phenotype. J Invest Dermatol. 2000;114(6): 1101–7.

[78] Niyonsaba F, Ushio H, Nakano N, Ng W, Sayama K, Hashimoto K et al. Antimicrobial peptides human beta-defensins stimulate epidermal keratinocyte migration, proliferation and production of proinflammatory cytokines and chemokines. J Invest Dermatol. 2007;127(3): 594–604.

[75] Rossini M, Viapiana O, Adami S, Idolazzi L, Fracassi E, Gatti D. Focal bone involvement in inflammatory arthritis: the role of IL17. Rheumatol Int. 2016;36(4): 469–82.

[76] Uluçan Ö, Jimenez M, Karbach S, Jeschke A, Graña O, Keller J, et al. Chronic skin inflammation leads to bone loss by IL-17-mediated inhibition of Wnt signaling in osteoblasts. Sci Transl Med. 2016;8(330): 330ra37.

[77] McGonagle D, Tan AL, Benjamin M. The nail as a musculoskeletal appendage - Implications for an improved understanding of the link between psoriasis and arthritis. Dermatology. 2009;218(2): 97–102.

[78] Pollitt CC. Anatomy and physiology of the inner hoof wall. Clin Tech Equine Pract. 2004;3: 3–21.

[79] Mosteller RD. Simplified calculation of body-surface area. N Engl J Med. 1987;317(17): 1098.

[80] Dern K, Burns TA, Watts MR, van Eps AW, Belknap JK. 2019. Influence of digital hypothermia on lamellar events related to IL-6/gp130 signalling in equine sepsis-related laminitis. Equine Vet J. 2020;52(3): 441–8.

[81] Lane HE, Burns TA, Hegedus OC, Watts MR, Weber PS, Woltman KA, et al. Lamellar events related to insulin-like growth factor-1 receptor signalling in two models relevant to endocrinopathic laminitis. Equine Vet J. 2017;49(5): 643–54.

[82] Ren W, Yin J, Duan J, Liu G, Tan B, Yang G, et al. mTORC1 signaling and IL-17 expression: Defining pathways and possible therapeutic targets. Eur J Immunol. 2016; 46(2): 291–9.

[83] Moss KL, Jiang Z, Dodson ME, Linardi RL, Haughan J, Gale AL, et al. Sustained interleukin-10 transgene expression following intra-articular AAV5-IL-10 administration to horses. Human Gene Ther. 2020;31(1-2): 110–18.

